# Gamma-irradiated SARS-CoV-2 vaccine candidate, OZG-38.61.3, confers protection from SARS-CoV-2 challenge in human ACEII-transgenic mice

**DOI:** 10.1101/2020.10.28.356667

**Authors:** Raife Dilek Turan, Cihan Tastan, Derya Dilek Kancagi, Bulut Yurtsever, Gozde Sir Karakus, Samed Ozer, Selen Abanuz, Didem Cakirsoy, Gamze Tumentemur, Sevda Demir, Utku Seyis, Recai Kuzay, Muhammer Elek, Miyase Ezgi Kocaoglu, Gurcan Ertop, Serap Arbak, Merve Acikel Elmas, Cansu Hemsinlioglu, Ozden Hatirnaz Ng, Sezer Akyoney, Ilayda Sahin, Cavit Kerem Kayhan, Fatma Tokat, Gurler Akpinar, Murat Kasap, Ayse Sesin Kocagoz, Ugur Ozbek, Dilek Telci, Fikrettin Sahin, Koray Yalcin, Siret Ratip, Umit Ince, Ercument Ovali

## Abstract

The SARS-CoV-2 virus caused the most severe pandemic around the world, and vaccine development for urgent use became a crucial issue. Inactivated virus formulated vaccines such as Hepatitis A, oral polio vaccine, and smallpox proved to be reliable approaches for immunization for prolonged periods. During the pandemic, we produced an inactivated SARS-CoV-2 vaccine candidate, having the advantages of being manufactured rapidly and tested easily in comparison with recombinant vaccines. In this study, an inactivated virus vaccine that includes a gamma irradiation process for the inactivation as an alternative to classical chemical inactivation methods so that there is no extra purification required has been optimized. The vaccine candidate (OZG-38.61.3) was then applied in mice by employing the intradermal route, which decreased the requirement of a higher concentration of inactivated virus for proper immunization, unlike most of the classical inactivated vaccine treatments. Hence, the novelty of our vaccine candidate (OZG-38.61.3) is that it is a non-adjuvant added, gamma-irradiated, and intradermally applied inactive viral vaccine. Efficiency and safety dose (either 10^13^ or 10^14^ viral copy per dose) of OZG-38.61.3 was initially determined in Balb/c mice. This was followed by testing the immunogenicity and protective efficacy of OZG-38.61.3. Human ACE2-encoding transgenic mice were immunized and then infected with a dose of infective SARS-CoV-2 virus for the challenge test. Findings of this study show that vaccinated mice have lower SARS-CoV-2 viral copy number in oropharyngeal specimens along with humoral and cellular immune responses against the SARS-CoV-2, including the neutralizing antibodies similar to those shown in Balb/c mice without substantial toxicity. Subsequently, plans are being made for the commencement of Phase 1 clinical trial of the OZG-38.61.3 vaccine for the COVID-19 pandemic.

## Introduction

The development of a vaccine has the upmost biomedical priority due to the global COVID-19 pandemic caused by the SARS-CoV-2 virus. A safe and effective SARS-CoV-2 vaccine is urgently required to halt the global COVID-19 pandemic. Several vaccine candidates have started clinical trials and some are in still preclinical research (Gao et al. 2020a; B. L. Corey et al., 2020; Yu et al. 2020). Small animal model systems are critical for better understanding the COVID-19 disease pathways and to determine medical precautions for improved global health, considering that there are currently no approved vaccines and only one antiviral approved for emergency use for SARS-CoV-2 (Sheahan et al. 2017; Dinnon et al. 2020). More significantly, several pioneering studies have shown that both SARS-CoV-2 and SARS-CoV use the same human angiotensin-converting enzyme 2 (hACE2) cellular receptor to enter cells (Walls et al. 2020; Li et al. 2003; Zhou et al. 2020a; Sun et al. 2020). The crystal structure of the SARS-CoV-2 S protein receptor-binding domain (RBD) which binds to hACE2 has been described, with an approximately 10- to 20-fold greater affinity toward hACE2 than SARS-CoV binds. Unfortunately, standard laboratory mice cannot be infected with SARS-CoV-2 due to the discrepancy of the S protein to the murine orthologous (mACE2) of the human receptor, making model development complicated and difficult (Zhou et al. 2020b; Dinnon et al. 2020). Thus, wild-type C57BL/6 mice cannot be infected efficiently with SARS-CoV-2 because there is no hACE2 protein expressed that supports SARS-CoV-2 binding and infection. On the other hand, both young and aged hACE2 positive mice showed high viral loads in the lung, trachea, and brain upon intranasal infection in the literature (Sun et al. 2020; Letko, Marzi, and Munster 2020; Wan et al. 2020; Winkler et al. 2020).

For understanding viral pathogenesis, vaccine production, and drug screening, animal models are crucial. To assess preclinical efficacy, non-human primates (NHPs) are the best animal models. The implementation of NHPs, however, is limited by the high costs, availability, and complexity of the necessary husbandry settings. For research and antiviral therapeutic progress, suitable small animal models are therefore important. Mouse models are popular because of their affordability, availability, and simple genetic structure, and have been commonly used to research human coronavirus pathogenesis (Cockrell et al. 2018; Jiang et al. 2020). As a cellular receptor, SARS-CoV-2 could use the ACE2 receptor of the human, bat, or civet but not the mouse (Jiang et al. 2020; Zhou et al. 2020b). Therefore, it seems that mice expressing hACE2 would be a conceivable choice for the vaccine challenge tests.

In this study, we tested our vaccine candidate (OZG-38.61.3) inactivated with gamma irradiation to assess their immunogenicity and protective efficacy against the SARS-CoV-2 viral challenge in K18-hACE2 mice and showed the efficacy of the vaccination in c57/Balb/C mice. K18-hACE2-transgenic mice, in which hACE2 expression is powered by the epithelial cell cytokeratin-18 (K18) promoter, were originally designed for the study of SARS-CoV pathogenesis and lead to a lethal infection model (McCray et al. 2007; Yang, Pabon, and Murry 2014; Winkler et al. 2020). This study aimed to investigate whether the vaccinated transgenic mouse has a lower SARS-CoV-2 viral copy number in nasal specimens along with increased humoral and cellular immune responses, including neutralizing antibodies to the virus, without experiencing substantial toxicity.

## Material & Methods

### Human Samples

In vitro isolation and propagation of SARS-CoV-2 from diagnosed COVID-19 patients were described in our previous study (Taştan et al., 2020). The study for SARS-CoV-2 genome sequencing was approved by the Ethics Committee of Acıbadem Mehmet Ali Aydınlar University (ATADEK-2020/05/41) and informed consent from the patients was obtained to publish identifying information/images. These data do not contain any private information of the patients. All techniques had been executed according to the applicable guidelines.

### Manufacturing Gamma-irradiated inactivated SARS-CoV-2 vaccine candidate

For the nasopharyngeal and oropharyngeal swab samples to have clinical significance, it is extremely important to comply with the rules regarding sample selection, taking into the appropriate transfer solution, transportation to the laboratory, and storage under appropriate conditions when necessary (Taştan et al., 2020). The production of a candidate vaccine for gamma-irradiated inactivated SARS-CoV-2 was reported in our previous report (Sir Karakus et al. 2021). In this study, the last version of our vaccine candidate, OZG-38.61.3 was constituted from 10^13^ or 10^14^ viral copy of SARS-CoV-2 in a dose without adjuvant.

### Viral RNA Extraction and Viral Genome Sequencing

Viral RNA extractions were performed by Quick-RNA Viral Kit (Zymo Research, USA) in Acıbadem Labcell Cellular Therapy Laboratory BSL-3 Unit according to the manufacturer’s protocols. Library preparation was performed by CleanPlex SARS-CoV-2 Research and Surveillance NGS Panel (Paragon Genomics, USA) according to the manufacturer’s user guide.

For the construction of the library, The CleanPlex® Dual-Indexed PCR Primers for Illumina® (Paragon Genomics, USA) were used by combining i5 and i7 primers. Samples were sequenced by Illumina MiSeq instrument with paired-end 131 bp long fragments. The data that passed the quality control were aligned to the reference genome (NC_045512.2) in Wuhan and a variant list was created with variant calling. The data analysis was described in detail in our previous study (Ozden Hatirnaz Ng et al., under revision).

### Nanosight

Nanoparticle Tracking Analysis (NTA) measurements were carried out for SARS-CoV-2 titer in suspension by using The NanoSight NS300 (Amesbury, UK). Samples were diluted with distilled water 1:10 ratio and transferred to Nanosight cuvette as 1 ml. Measurements were performed at room temperature with 5 different 60-second video recording.

### Inactivated SARS-CoV-2 virus imaging by transmission electron microscopy

Viruses were inactivated and fixed with 2.5% glutaraldehyde in PBS (0.1 M, pH 7.2) for 2.5 h. One drop of glutaraldehyde-treated virus suspension was placed on the carbon-coated grid for 10 min. The remaining solution was absorbed with a filter paper and the grid was stained by a negative staining procedure. Then, it was evaluated under a transmission electron microscope (Thermo Fisher Scientific-Talos L120C) and photographed.

### In-solution tryptic digestion

In-solution digestion was performed according to the manufacturer’s instructions using ‘in-solution tryptic digestion and guanidination kit’ (#89895, Thermo Fisher Scientific, USA). The protocol can be summarized as follow: 10 μg protein sample was added to 15 μL 50 mM Ambic containing 100 mM DTT solution. The volume was completed to 27 μL and incubated at 95°C for 5 min. Iodoacetamide (IAA) was added to the heated sample to a 10 mM final concentration and incubated in the dark for 20 min. 1 μL of 100ng/μL trypsin was then added and incubated for 3 hours at 37°C. 1 μL of 100ng/μL trypsin was added to the peptide mixture and incubated overnight at 30°C. After incubation, the solution was vacuum concentrated to dryness and the peptides were resuspended in 0.1% FA for the nLC-MS/MS analysis.

### Nano-liquid chromatography mass spectrometry (nLC-MS/MS) analysis

The peptides were analyzed by nLC-MS/MS using an Ultimate 3000 RSLC nanosystem (Dionex, Thermo Scientific, USA) coupled to a Q Exactive mass spectrometer (Thermo Scientific, USA). The entire system was controlled by Xcalibur 4.0 software (Thermo Fisher Scientific, USA). High-performance liquid chromatography(HPLC) separation was performed using mobiles phases of A (%0.1 Formic Acid) and B (%80 Acetonitril+%0.1 Formic Acid). Digested peptides were pre-concentrated and desalted on a trap column. Then the peptides were transferred to an Acclaim PepMap RSLC C18 analytical column (75 μmx15 cmx2 μm, 100 Å diameter, Thermo Scientific, USA). The gradient for separation was 6-32% B in 80 min, 32-50% B in 40 min, 50-90% B in 10 min, 90% in 15 min, 90-6% B in 10 min, and 6% B for 10 min with the flow rate of 300 nL/min. Full scan MS spectra were acquired with the following parameters: resolution 70.000, scan range 400-2000 m/z, target automatic gain control (AGC)3×106, maximum injection time 60 ms, spray voltage 2.3 kV. MS/MS analysis was performed by data-dependent acquisition selecting the top ten precursor ions. The instrument was calibrated using a standard positive calibrant (LTQ Velos ESI Positive Ion Calibration Solution 88323, Pierce, USA) before each analysis.

### nLC-MS/MS data analysis

Raw data were analyzed with Proteom Discoverer 2.2 (Thermo Scientific, USA) software for protein identification and the following parameters were used; peptide mass tolerance 10 ppm, MS/MS mass tolerance 0.2 Da, mass accuracy 2 ppm, tolerant miscarriage 1, minimum peptide length 6 amino acids, fixed changes cysteine carbamidomethylation, unstable changes methionine oxidation, and asparagine deamination. The minimum number of peptides identified for each protein was considered to be 1 and obtained data were searched in the Uniprot/Swissprot database.

### Vero Host Cell Protein ELISA

Residual Host Cell Protein (HCP) analysis in a viral product supernatant was performed with the manufacturer’s protocol of the Cygnustechnologies-VERO Cell HCP ELISA kit (F500). The absorbance was read at 450/650nm with the microplate reader (Omega ELISA Reader).

### Vero DNA nanodrop

The vaccine candidate was solved in 100 cc pyrogen-free water. Firstly, pyrogen-free water was blanked and one drop sample was measured at the dsDNA program using Thermo Scientific NanoDrop™ One Spectrophotometers to determine Vero residual DNA and A260/A280 ratio for DNA/protein purity.

### Replicative Competent Coronavirus test with gamma-irradiated inactivated SARS-CoV-2 vaccine candidates

3µg of lyophilized inactivated SARS-CoV-2 vaccine candidate in 100 µl pyrogen-free water was inoculated into %90 confluent Vero cells at 37C. The supernatant of this culture was replenished with fresh Vero cell culture every 3-to-5 days up to 21 days of incubation. As a negative control, only 100 µl pyrogen-free water was inoculated into Vero cells and cultured for 21 days with the same treatments. At the end of the incubation, the final supernatant was collected, centrifuged at 2000G for 10 min to remove cell debris. Next, the supernatants were concentrated 10x with 100kDa Amplicon tubes. The concentrated samples were tested in the xCelligence RTCA system in a dose-dependent manner as 10-1 to 10-6 to determine the cytopathic effect.

### SRID assay

5 μg / ml of SARS-COV-2 Spike S1 Monoclonal Antibody (ElabScience) antibodies were added in the gel at a concentration of 2%. Inactive SARS-CoV-2 was kept at room temperature for 15-30 minutes with 1% zwittergent detergent (mix 9 test antigens: 1 Zwittergent). Incubation was provided in a humid environment for 18 hours. The gel was washed with PBS, taken on the glass surface, and covered with blotter paper, and kept at 37 ° C until it dried. By staining the gel with Coomassie Brillant Blue, the presence of S antigen was determined according to the dark blue color (colorimetric).

### Quality Control Tests

Sterility was done in a BACTEC blood culture bottle along with the BACTEC™ FX blood culturing instrument (BD). The endotoxin level was determined with the Gel-clot endotoxin Limulus Amebocyte Lysate (LAL) test (Charles River Laboratories). Mycoplasma analysis was performed with Mycoplasma species 500 PCR kit at GeneAmp PCR System 2700 (Applied Biosystems). Quality control tests of the vaccine including levels of chemistry analysis (Na, Cl, K, Ca) and Total Protein (The ADVIA 1800 Clinical Chemistry System, Siemens), osmolarity (Osmometer, freezing point depression), Ph, Glucose, Albumin (Dimension EX-L), sterility, mycoplasma, endotoxin level, and impurity assay were performed in Acıbadem Labmed Laboratory with accredited methods. Moisture Analyzer was performed at Yeditepe University with accredited methods.

### Quantitative RT-PCR to determine viral copy number

Total RNA isolations were performed from SARS-CoV-2 specimens using Direct-zol RNA Miniprep Kits (Zymo Research, USA). Quantitative RT-PCR was performed with the QuantiVirus SARS-CoV-2 Test Kit (Diacarta) according to the manufacturer’s protocol. The quantitative RT-PCR analysis was analyzed in Roche Lightcycler 96.

### Balb/c mice test

To analyze the efficiency and toxicology of the dose of inactive vaccine candidate parallel to challenge, 15 Female BALB/c mice were utilized from Acıbadem Mehmet Ali Aydinlar University Laboratory Animal Application and Research Center (ACUDEHAM; Istanbul, Turkey). All animal studies received ethical approval by the Acibadem Mehmet Ali Aydinlar University Animal Experiments Local Ethics Committee (ACU-HADYEK). BALB/c mice were randomly allocated into 3 groups, a negative control group (n=5) and 2 different dose groups (dose of 1×10^13^ and 1×10^14^, n = 5 per group). To determine the immunogenicity with two different doses (dose 10^13^ and dose 10^14^, n=5 per group) of inactive vaccine produced in Acibadem Labcell Cellular Therapy Laboratory, Istanbul, Turkey, on day 0 mice were vaccinated intradermally with the dose of 1×10^13^ and 1×10^14^ lyophilized vaccine candidate without adjuvant reconstituted in 100 cc pyrogen-free water and also control groups vaccinated with 100 cc pyrogen-free water. After 18 days booster dose was applied with the same vaccination strategies. Survival and weight change were evaluated daily and every week respectively. Blood samples were collected just before the sacrification on day 28 for serum preparation to be used for in vitro efficiency studies. Mice were sacrificed on day 28 post-immunization for analysis of B and T cell immune responses via SARS-Cov-2 specific IgG ELISA, IFNγ ELISPOT, and cytokine bead array analysis. Furthermore, dissected organs including the lungs, liver, kidneys of sacrificed mice were taken into 10% buffered formalin solution before they were got routine tissue processing for histopathological analysis. Also, the spleen tissues were taken into a normal saline solution including %2 Pen-Strep for T cell isolation following homogenization protocol.

### Transgenic mice for Challenge test

5 female and 20 male B6.Cg-Tg(K18-hACE2)2Prlmn/J transgenic mice at 6 weeks of age were purchased from The Jackson laboratories. All animal experiments were approved by the Experimental Animal Committee of Acıbadem Mehmet Ali Aydınlar University (ACUHADYEK 2020/36). The mice housed in Transgenic Biosafety BSL-3 laboratories of AAALAC International accredited Acıbadem Mehmet Ali Aydinlar University Laboratory Animal Application and Research Center (ACUDEHAM; Istanbul, Turkey). Light, temperature, humidity, and feeding conditions followed the ACUDEHAM accredited operating procedures and also K18-hACE2 mice hospitalized in IVC systems (ZOONLAB BIO. A.S.) for 29-day challenge tests. Whole groups were identified as female and male in the base of the earring numbers start 40 to 64.

### Vaccination and Challenge Strategies

Transgenic mice were randomly allocated into 4 groups, negative control group (n=5), positive control group (n=6), and 2 different dose groups (dose of 1×10^13^ and 1×10^14^, n = 7 per group). To determine the 21-day immunogenicity with two different doses (dose 10^13^ and dose 10^14^, n=7 per group) of inactive vaccine produced in Acibadem Labcell Cellular Therapy Laboratory, Istanbul, Turkey, on day 0 mice were vaccinated intradermally with the dose of 1×10^13^ and 1×10^14^ SARS-CoV-2 viral copy per microliter lyophilized vaccine without adjuvant reconstituted in 100 cc pyrogen-free water and both negative and positive control groups vaccinated with 100 cc pyrogen-free water. In whole groups, a booster dose of 1×10^13^ and 1×10^14^ SARS-CoV-2 viral copy per microliter vaccine was administered intradermally on day 15 post-first vaccination. All animals were monitored daily for clinical symptoms, body-weight changes body temperature change (**Supplementary figure 1**). 25 days following vaccination, K18-hACE2 mice were intranasally infected with a 3×10^4^ TCID50 dose of infective SARS-CoV-2 in 30µl solution in Biosafety level cabin II in Transgenic Animal Biosafety level 3 laboratory (ABSL-3) of AAALAC International accredited Acıbadem Mehmet Ali Aydinlar University Laboratory Animal Application and Research Center (ACUDEHAM; Istanbul, Turkey). TCID50 dose of SARS-CoV-2 was calculated in the previous study (Sir Karakus et al. 2021). Starting from the day after the challenge, clinical symptoms, body-weight changes body temperature change controlled every 12 hours. At 48 hours after the challenge, the oropharyngeal swabs were collected from mice in all groups and analyzed for viral copy number. At 96 hours after the challenge, the nasopharyngeal swabs and sera were collected from whole groups including negative control groups to analyze immunological and virological assays. After serum collection, all mice were euthanized. Biopsy samples were collected including skin which was the vaccination part, brain, testis, ovarium, intestine, spleen, kidney, liver, lung, heart. Biopsy samples were collected and anatomically divided for qPCR analysis and histological and TEM examination.

**Figure 1.**
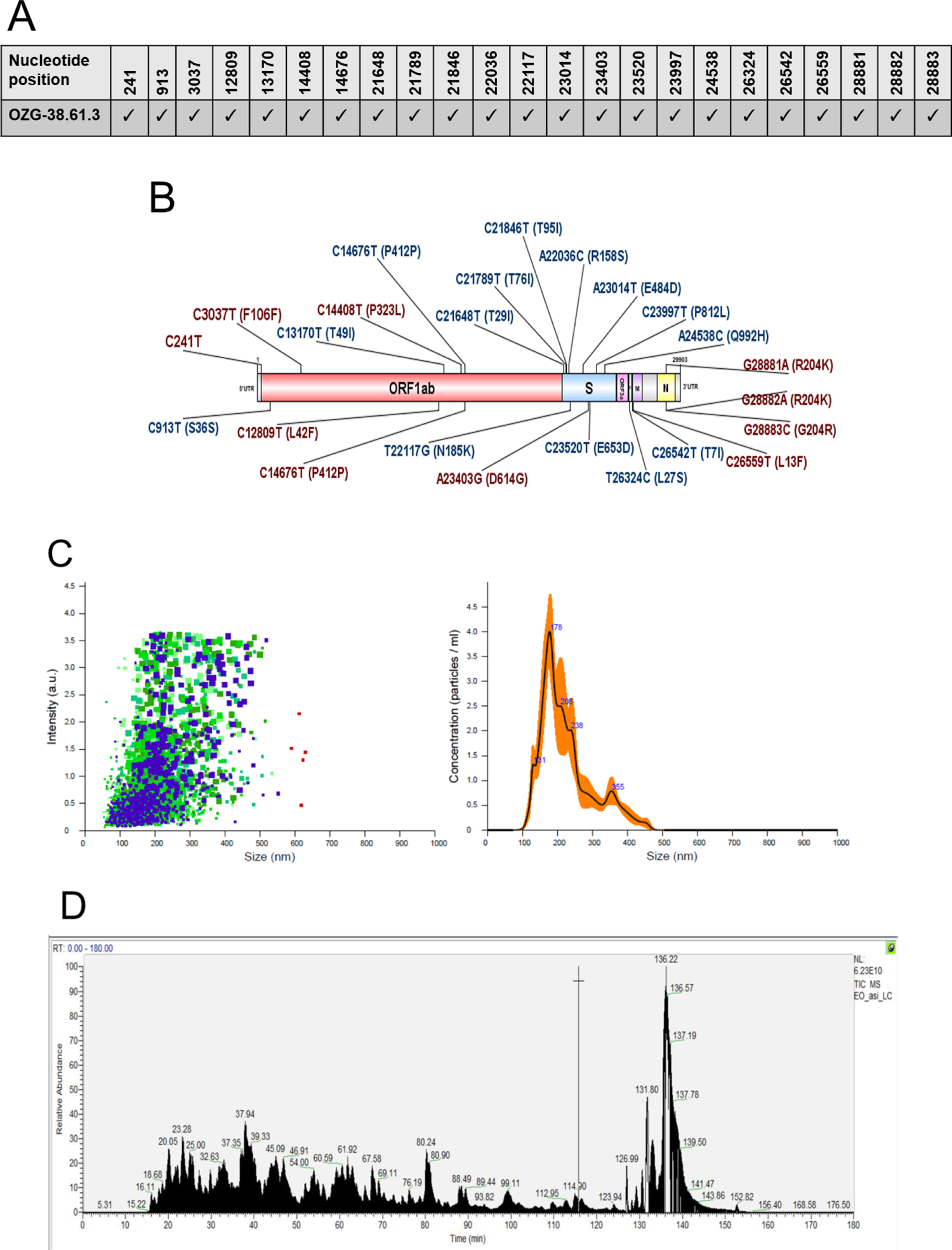
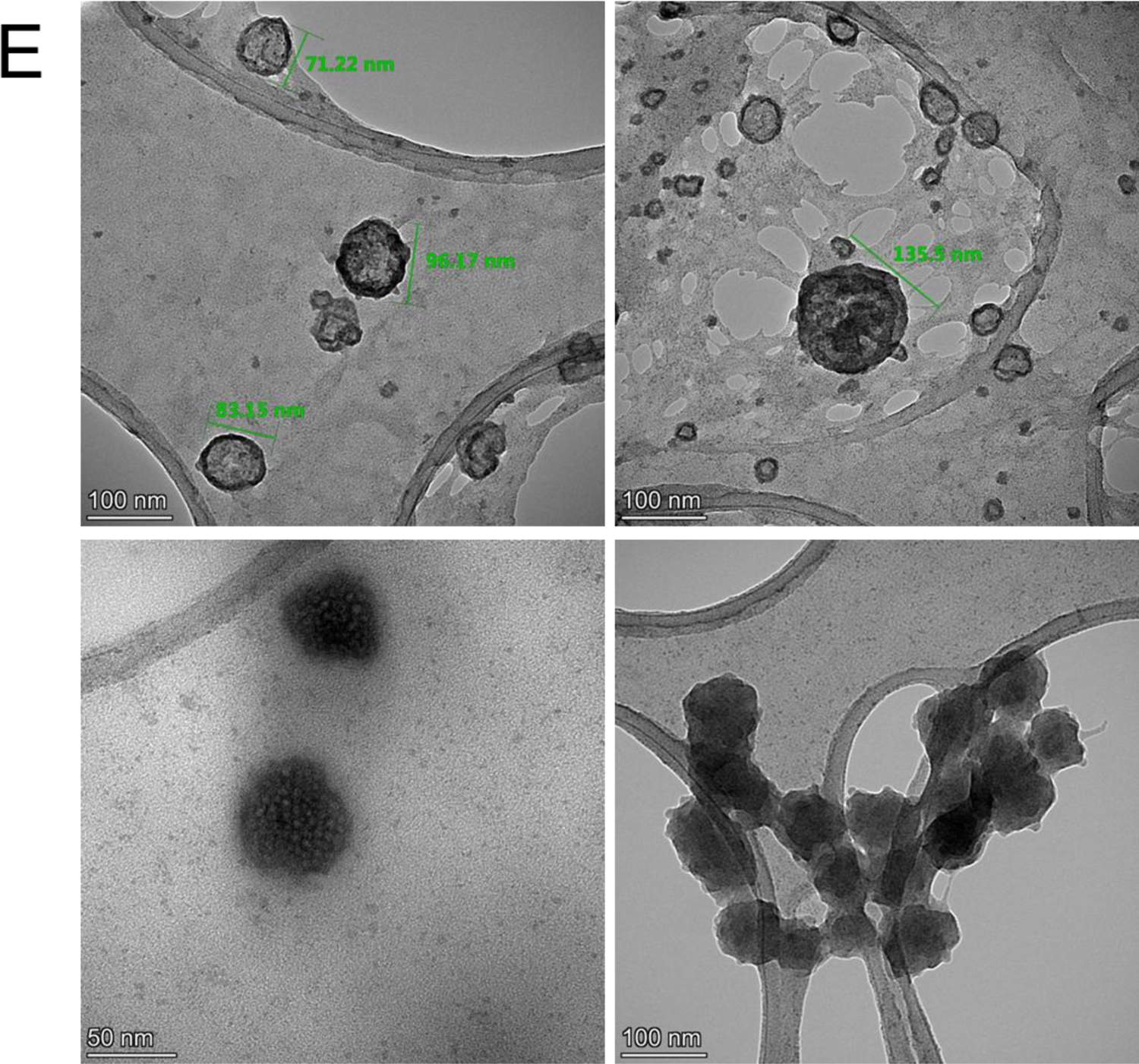
Characterization of inactivated SARS-CoV-2 virus constituting OZG-38.61.3 vaccine candidates. **A.** Mutation distribution of SARS-CoV-2 virus strain that makes up the OZG-38.61.3 vaccine candidate. **B.** Representation of variants detected in the virus strain that makes up the OZG-38.61.3 vaccine candidate on the SARS-CoV-2 genome. **C.** The left plot showing intensity versus the size of the particles in OZG-38.61.3. The right plot showing means of particle size of the candidate in the sample read three times. **D.** Proteome analysis of inactivated OZG-38.61.3 SARS-CoV-2 product. **E.** TEM Image of SARS-CoV-2 Virus. Representative electron micrographs of SARS-CoV-2. Virus particles were seen on the grid (Scale bars: 50 nm, 100 nm).

### X-ray dark-field Imaging of the Lungs of SARS-CoV-2−Infected K18-hACE2 mice

At 96 hours after the challenge, whole mice of each group imaged with the Siemens Arcadis Avantic C arms X-ray dark-field imaging system to evaluate the feasibility of early-stage imaging of acute lung inflammation in mice. 3 mice from each group were imaged and also all mice anesthetized once during imaging with Matrx VIP 3000 Isoflurane Vaporizer (MIDMARK) system. All images were acquired as the posterior prone position of mice. The X-ray ran at 48 kV, distance to source grating 70cm, 111°, and shooting with 0.2 and 0.3 mA.

### Histopathological Applications

Transgenic mice and Balb/c mice were sacrificed on postimmunization for histopathology analysis. Dissected organs including the cerebellum, lungs, liver, kidneys, skin, intestine, and part of the spleen of sacrificed mice were taken into 10% buffered formalin solution before routine tissue processing for histopathological analysis after weighting. The histopathology analysis of the lung tissues of challenge mice groups was performed at the Department of Pathology at Acibadem Maslak Hospital.

### SARS-CoV-2 IgG ELISA

Before the sacrification, blood samples were collected from the whole group of mice. The serum was collected with centrifugation methods. Serum samples were stored at −40 C. To detect the SARS-COV-2 IgG antibody in mouse serum SARS-CoV-2 IgG ELISA Kit (Creative, DEIASL019) was used. Before starting the experiment with the whole sample, reagent and microplates pre-coated with whole SARS-CoV-2 lysate were brought to room temperature. As a positive control, 100 ng mouse SARS-CoV-2 Spike S1 monoclonal antibody was used (commercially available as E-AB-V1005, Elabscience). Serum samples were diluted at 1:64, 1:128, and 1:256 in a sample diluent, provided in the kit. Anti-mouse IgG conjugated with Horseradish peroxidase enzyme (mHRP enzyme) was used as a detector. After incubation with the stopping solution, the color change was read at 450nm with the microplate reader (Omega ELISA Reader).

### Neutralization assay using colorimetric MTT assay

TCID50 (Median Tissue Culture Infectious Dose) of SARS-CoV-2 was determined by incubating the virus in a serial dilution manner with the Vero cell line (CCL81, ATCC) in gold microelectrodes embedded microtiter wells in xCELLigence Real-Time Cell Analysis (RTCA) instruments (ACEA, Roche) for 8 days (Sir Karakus et al. 2021). Neutralization assay of sera from transgenic and balb/c mice groups was performed at 1:128, and 1:256 dilutions pre-incubated with a 100X TCID50 dose of SARS-CoV-2 at room temperature for 60 min. Next, the pre-incubated mixture was inoculated into the Vero-cell-coated flat-bottom 96-well plate which was analyzed at the end of 96 hr following standard MTT protocol. Viable cell analysis was determined by colorimetric change at the ELISA system. The neutralization ratio was determined by assessing percent neutralization by dividing the value of serum-virus treated condition wells by the value of untreated control Vero cells. 100% of neutralization was normalized to only Vero condition while 0% of neutralization was normalized to the value of only 100x TCID50 dose of SARS-CoV-2 inoculated Vero cell condition. For example, for the sample of 1:128 serum sample, the value was 0,651 while the value for control Vero well as 0,715, and the value for control SARS-CoV-2 inoculated well was 0,2. The calculation is as %neutralization= ((0,651-0,2)*100)/(0,715-0,2). This gave 87,5% virus neutralization. This calculation was performed for each mouse in the group and the mean of the virus neutralization was determined.

### Mouse IFN-γ ELISPOT analysis

Mouse Spleen T cells were centrifuged with Phosphate Buffer Saline (PBS) at 300xg for 10 min. Pellet was resuspended in TexMACs (Miltenyi Biotech, GmbH, Bergisch Gladbach, Germany) cell culture media (%3 human AB serum and 1% Pen/Strep). 500,000 cells in 100 µl were added into microplate already coated with a monoclonal antibody specific for mouse IFN-γ. 1000 nM SARS-CoV-2 virus Peptivator pool (SARS-CoV-2 S, N, and M protein peptide pool) (Miltenyi Biotech, GmbH, Bergisch Gladbach, Germany) were added into each well including mouse spleen T cells. The microplate was incubated in a humidified 37°C CO_2_ incubator. After 48 h incubation, IFN-γ secreting cells were determined with Mouse IFNγ ELISpot Kit (RnDSystems, USA) according to the manufacturer’s instructions. The spots were counted under the dissection microscope (Zeiss, Germany).

### Unstimulated/Stimulated T cell cytokine response and immunophenotype

500,000 cells isolated from mouse spleen were incubated with 1000 nM SARS-CoV-2 virus Peptivator pool (SARS-CoV-2 S, N, and M protein peptide pool) (Miltenyi Biotech, GmbH, Bergisch Gladbach, Germany) in a humidified 37°C CO_2_ incubator. After 48h incubation, the mouse cytokine profile was analyzed using the supernatant of the cultures using the MACSPlex Cytokine 10 kit (Miltenyi Biotec). Also, to determine T cell activation and proliferation, the restimulated cells were stained with the antibodies including CD3, CD4, CD8, and CD25 as an activation marker (Miltenyi Biotec). The Cytokine bead array and the T cell activation and proportions were analyzed using the MACSQuant Analyzer (Miltenyi Biotec).

### Statistics

Normally distributed data in bar graphs was tested using student’s t-tests for two independent means. The Mann-Whitney U test was employed for comparison between two groups of non-normally distributed data. Statistical analyses were performed using the Graphpad Prism and SPSS Statistics software. Each data point represents an independent measurement. Bar plots report the mean and standard deviation of the mean. The threshold of significance for all tests was set at *p<0.05. *ns* is non-significant.

## Results

### Characterization of inactivated SARS-CoV-2 virus constituting OZG-38.61.3 vaccine candidates

One of the conclusions of our previous study was that the adjuvant positive vaccine administration should be removed from the newly designed version of the OZG-38.61 vaccine model as it caused inflammatory reaction in the skin, cerebellum, and kidney in toxicity analysis of vaccinated mice (Sir Karakus et al. 2021). Hence, it was decided to increase the SARS-CoV-2 effective viral copy dose (1×10^13^ or 1×10^14^ viral copies per dose) without an adjuvant in this new version of the OZG-38.61 vaccine. Firstly, we determined whether this vaccine product comprises all identified SARS-CoV-2 mutations. The obtained sequences from the propagated SARS-CoV-2 virus were compared with the GISAID database and the protein levels of the variant information were examined (**Fig. 1A**). We determined that the SARS-CoV-2 strain forming OZG-38.61.3 vaccine covered previously identified mutation variants (red-colored) with new variants (blue colored) (**Fig. 1B**). The data of all defined mutations were presented in detail in **Supplementary Table 2**. Using Nanosight technology, we determined that the size of inactivated SARS-CoV-2 was 187.9 +/- 10.0 nm (mode) with a concentration of 4.23×10^9^ +/- 1.88×10^8^ particles/ ml (**Fig. 1C**). As a result of the LC-MS-MS analysis, the presence of proteins belonging to the SARS-CoV-2 virus was detected in the analyzed sample (**Fig. 1D**). Quality control tests are illustrated in **Supplementary Table 3**. Four of the defined proteins were Master Proteins and have been identified with high reliability (**Supplementary Table 3**). Other proteins were Master Candidate proteins and their identification confidence interval is medium. The data of all defined proteins were presented in detail in **Supplementary Table 3**. Also, the transmission electron microscope was evaluated with the negative staining method and the main structures (envelope/spike) of the SARS-CoV-2 virus particles in the final product were well preserved (**Fig. 1E**). Also, TEM analysis confirmed the virus size was 70-200 nm with the presence of aggregates as determined in the Nanosight analysis. On the other hand, the concentration of the Vero host cell protein per vaccine dose was determined <4 ng and Vero host DNA per dose was not determined. According to all these analyses, OZG-38.61.3 has been shown to pass all vaccine development criteria in the final product (**Supplementary Table 1**), comprise all the mutations identified up to date, preserve the protein structure and contain pure inactive virus free of residues.

### Acute Toxicity and Efficacy Study of OZG-38.61.3 in Balb/c mice

Acute toxicity and efficacy assays were studied in Balb/c mice. There were 3 groups in this study: control group (n=5), dose 10^13^ viral particles per dose (n=5), and dose 10^14^ viral particles per dose (n=5) (**Fig. 2A**). There was no difference in nutrition and water consumption between the groups, as well as in total body weight and organ weight (**Supplementary Table 4**). Version 3 of the OZG-38.61 vaccine did not differ in the histopathologic analysis from the control group, including the highest dose of 10^14^ (**Supplementary Table 4**). Firstly, SARS-CoV-2 specific IgG at 1:128 dilution of serum isolated from mice groups showed a significant increase in the highest dose (1014) vaccinated group in comparison with the control group (**Fig. 2B**). Secondly, in order to determine the neutralization capacity of serum collected from the immunized mice, 1:128 and 1:256 dilutions of sera were pre-inoculated with the SARS-CoV-2 virus, and following 96-hour incubation, MTT analysis was performed. Findings showed that the dose 10^14^ vaccinated mice managed to neutralize the virus at a statistically significant level (p<0.05) at both dilutions (**Fig. 2C**). The Balb/c study showed that upon restimulation, gamma interferon secretion of the T lymphocytes from both vaccination groups increased significantly in comparison with the non-vaccinated mice group and PBS non-stimulated internal control groups (**Fig. 2D**). Furthermore, spleen T cells stimulated with the peptides were analyzed through flow cytometry in order to determine proportions of activated (CD25+) CD4+ and CD8+ T cells, but no proliferation was observed (**Fig. 2E**). Next, the supernatant of the incubated cells was analyzed using a cytokine bead array for a more detailed examination. Both doses of OZG-38.61.3 (especially dose 10^14^) increased IL-2, GM-CSF, gamma-IFN levels and caused Th-1 response (**Fig. 2F**). At the same time, IL-10 was increased in both dose groups, suggesting that the Tr1 (Regulatory T lymphocyte type 1 response) response was stimulated (**Fig. 2F**) (Andolfi et al. 2012). Moreover, we performed histopathology analysis of the lung, liver, and kidney to determine inflammation, hemorrhage, and eosinophil infiltration (**Fig. 2G** and **Supplementary Table 4**). There was no significant toxicity including hemorrhage and eosinophil infiltration in overall organs (**Fig. 2G** and **Supplementary Table 4**). Therefore, 1×10^13^ and 1×10^14^ doses of OZG-38.61.3 led to the effective neutralizing SARS-CoV-2 specific IgG antibody production and cytokine secretion, hence a satisfactory Th1 response, without significant toxicity.

**Figure 2:**
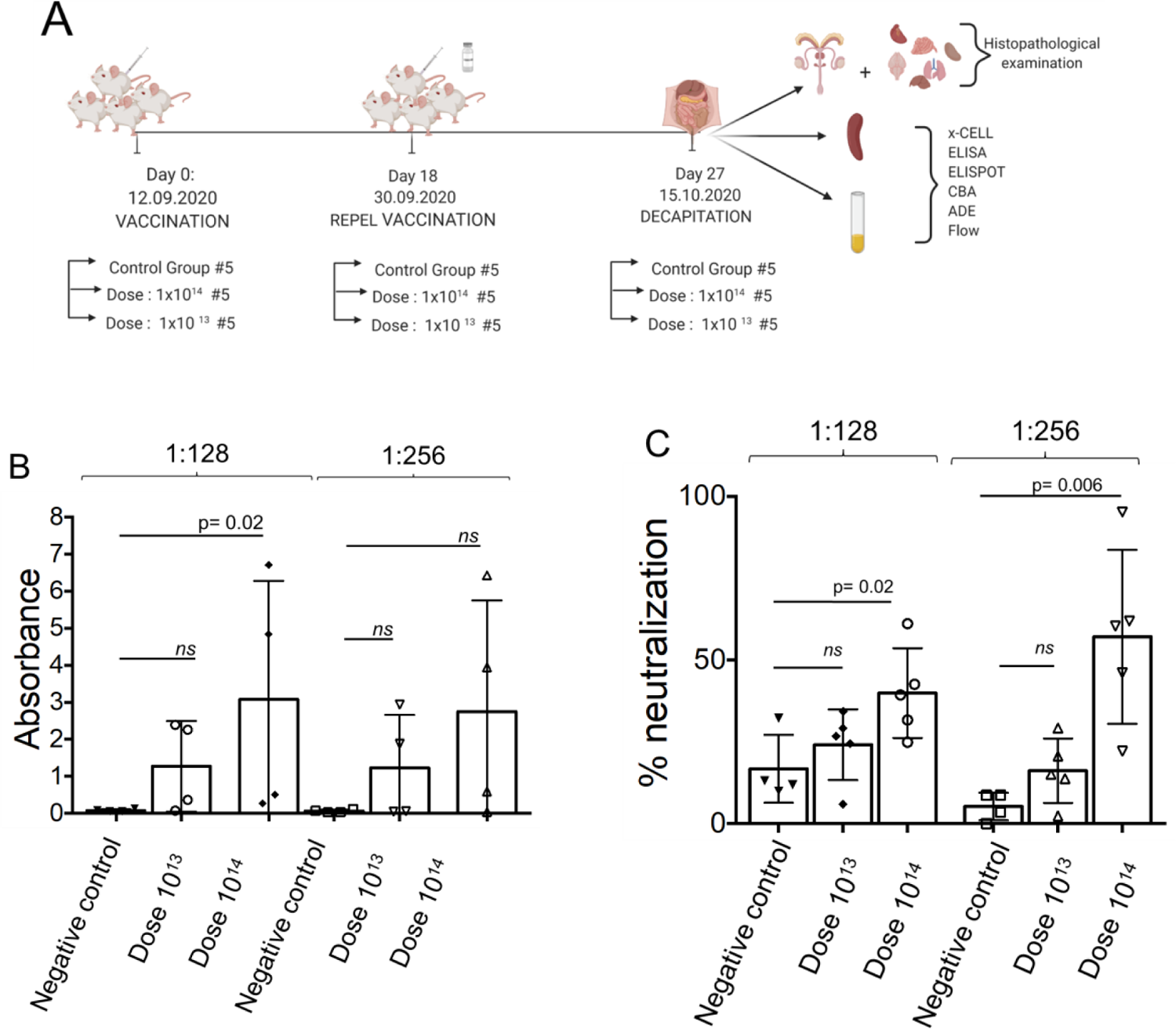
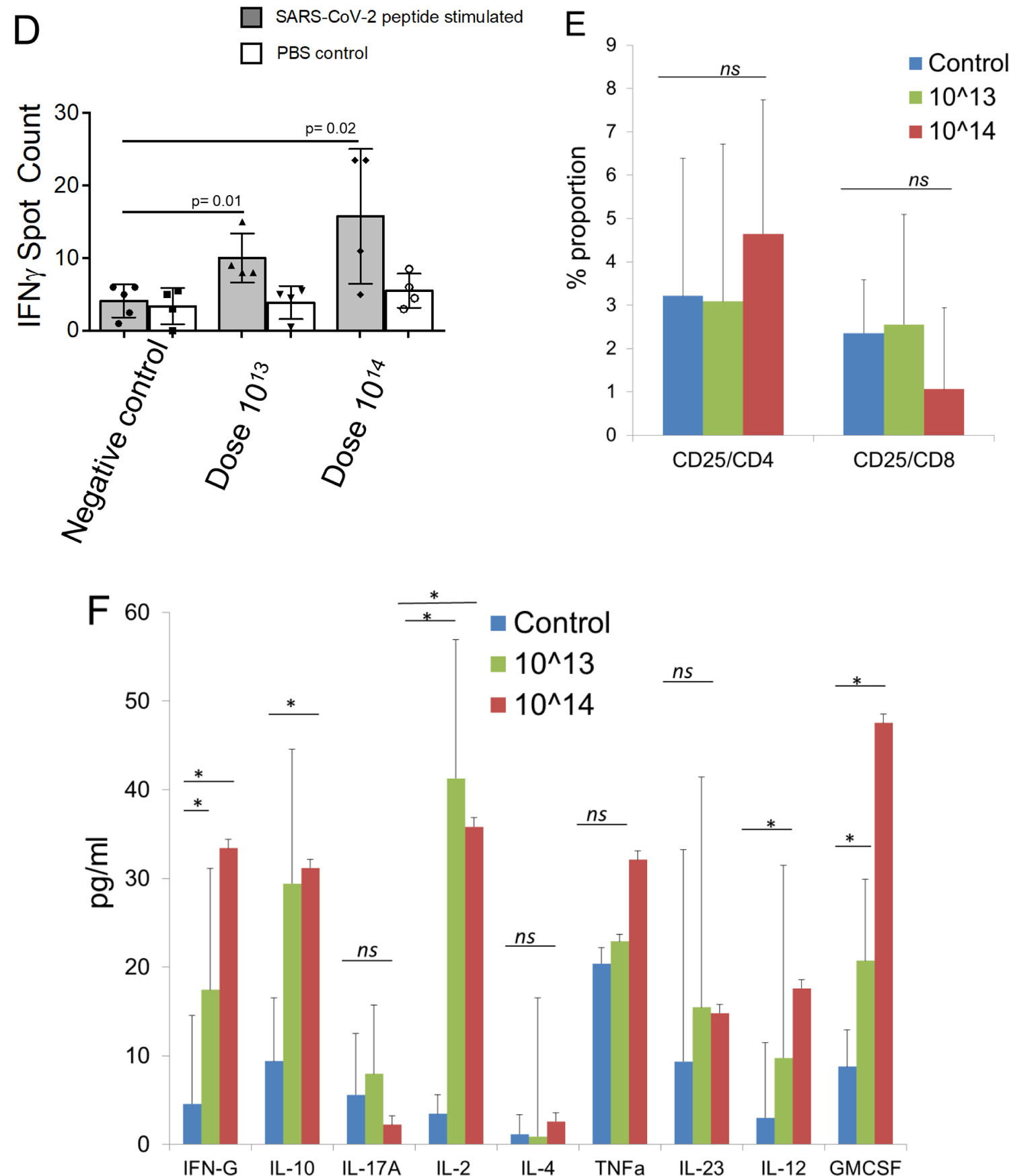
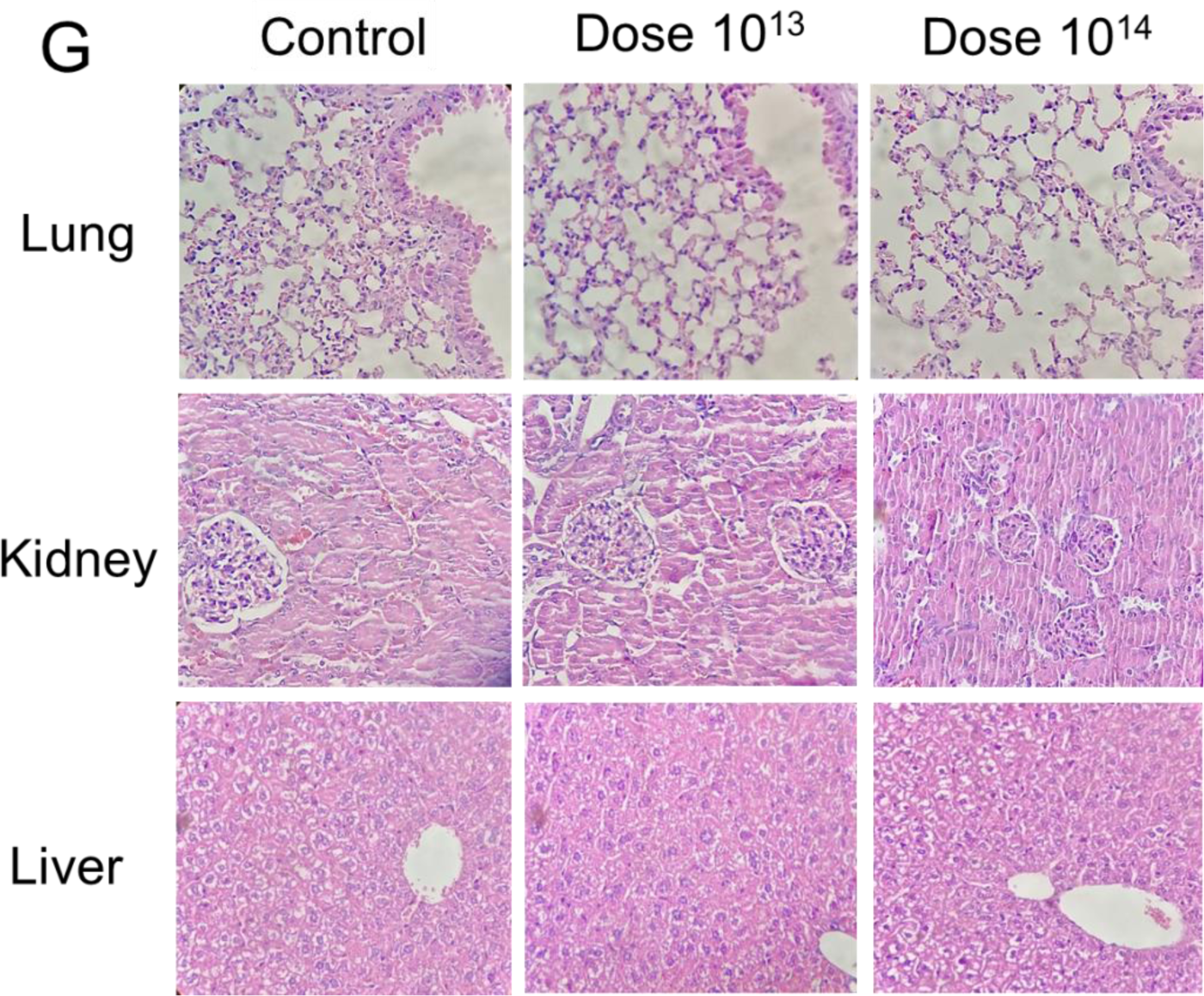
In vivo efficacy analysis of Dose 10^13^ and Dose 10^14^ in vaccinated Balb/c mice. **A.** Representation of in vivo experimental setup of the Balb/c mice vaccinated with Dose 10^13^ or Dose 10^14^ of OZG-38.61.3. **B.** The bar graph showing presence (absorbance) of SARS-CoV-2 specific IgG in the mice sera diluted with either 1:128 or 1:256 detected using ELISA. **C.** The bar graph showing neutralization frequency of the mice sera diluted to 1:128 or 1:256 that were pre-incubated with a 100x TCID50 dose of infective SARS-CoV-2. The analysis was performed with MTT analysis at 96hr. **D.** The bar graph showing activation frequency (IFNγ positive spot count) of spleen T cells that were incubated with SARS-CoV-2 specific peptides. **E.** The bar graph showing the proportion of the activated T cells after the stimulation. The activation was determined with the upregulation of the CD25 surface marker. **F**. The bar graph showing cytokine proportions of spleen T cells that were incubated with SARS-CoV-2 specific peptides. **G.** Histopathologic analysis of the lung, kidney, and liver tissues of Balb/c mice groups. H&E stain X400.

### Challenge Test with OZG-38.61.3 vaccinated humanized ACEII+ mice

Following efficacy and safety analysis of OZG-38.61.3 in Balb/c mice, post-immunization protection from SARS-CoV-2 infection in human ACE2 expressing transgenic mice was determined. Viral challenge analyzes were performed in K18-hACE2 (Jackson Lab). Two mice groups were vaccinated with the 10^13^ and 10^14^ doses of the vaccine. Mice were euthanized for in vitro efficacy tests and histopathology analysis post-challenge on day 4 (**Fig. 3A**). During the challenge, there was no significant change in food and water consumption along with temperature (**Supplementary Figure 1**). Also, although not statistically significant, weight distribution was more uniform in vaccine groups (**Supplementary Figure 1**).

**Figure 3.**
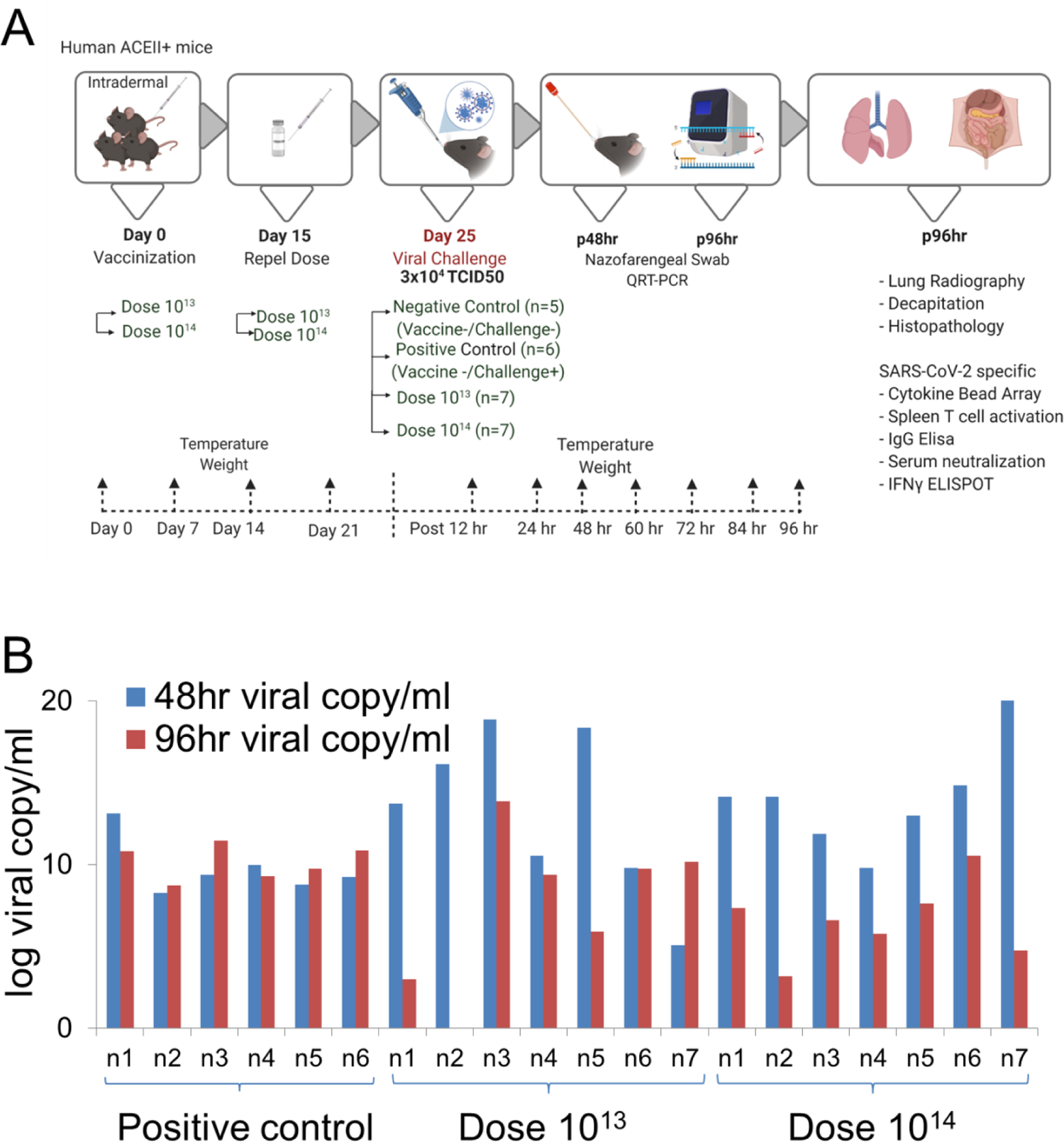
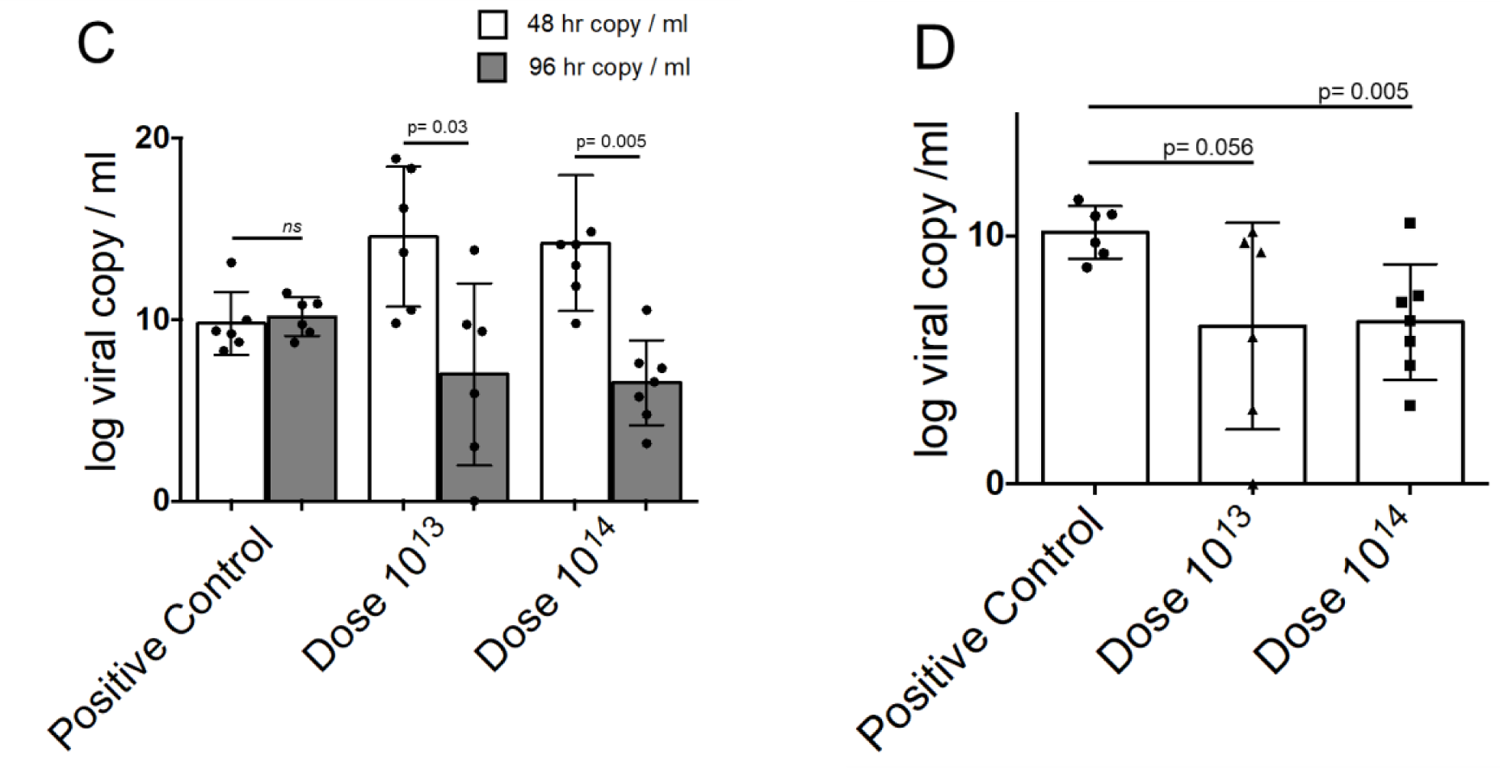
Challenge Test with OZG-38.61.3 vaccinated humanized ACEII+ mice. **A.** Representation of in vivo experimental setup of the challenge test. **A.** Mice were allocated into 4 groups, negative control group (n = 5), positive control group (n=6), and 2 different intradermally vaccinated group (dose 10^13^and dose 10^14^, n = 7 per group). A booster dose of dose 10^13^and dose 10^14^ vaccine was administered on day 15. After 25 days of vaccination, the mice were intranasally infected with a 3×10^4^ TCID50 dose of SARS-CoV-2. Biopsy samples, spleen T cells, and serum were collected after euthanization at 96hr. **B.** The bar graph showing SARS-CoV-2 viral copy number in log scale per ml of the nasopharyngeal samples collected from each mouse at 48hr and 96 hr post-challenge that were either vaccinated with dose 10^13^ (n=7), dose 10^14^ (n=7) or without vaccination group (positive control; n=6). **C.** The bar graph shows the mean value of viral copy in log scale per ml at 48hr and 96 hr post-challenge. **D.** The bar graph showing a comparison of viral copy number at 96hr between vaccinated and positive control groups.

This was followed by viral load analyzes on oropharyngeal swab samples on the 2nd and 4th days of the challenge test. While there was no significant change in the virus load in the positive control group, it was observed that the virus load decreased in the vaccine groups with the exception of only one mouse, whereas it completely disappeared in one (**Fig. 3B**). It was observed that the mean virus load decreased statistically significantly over time, especially at the highest dose (10^14^), between 48-96 hours. There was no change in the positive control group (**Fig. 3C**). When compared with the positive control group at 96th hour, a 3-log decrease in viral copy number was determined especially in the highest dose vaccine group (**Fig. 3D**). No difference was observed between the groups in the lung X-ray imaging analysis of the mice groups taken in our study (**Fig. 4A**). Also, it is observed that both vaccine doses do not cause antibody-dependent enhancement (ADE) side effects in the lung in histopathology analysis (**Fig. 4B**). No histologically significant change was observed in the positive control and vaccine groups, although positive control has signs of partial alveolar fusion and inflammation in 1 mouse (**Fig. 4B**). This finding is similar to chest radiographs. The absence of an additional pathology in the lung, especially in vaccine groups, was another additional finding confirming that ADE does not occur and the inactivity of our vaccine. Thus, viral load analyses in the oropharyngeal specimens showed that the SARS-CoV-2 infection was significantly reduced in the vaccinated groups.

**Figure 4.**
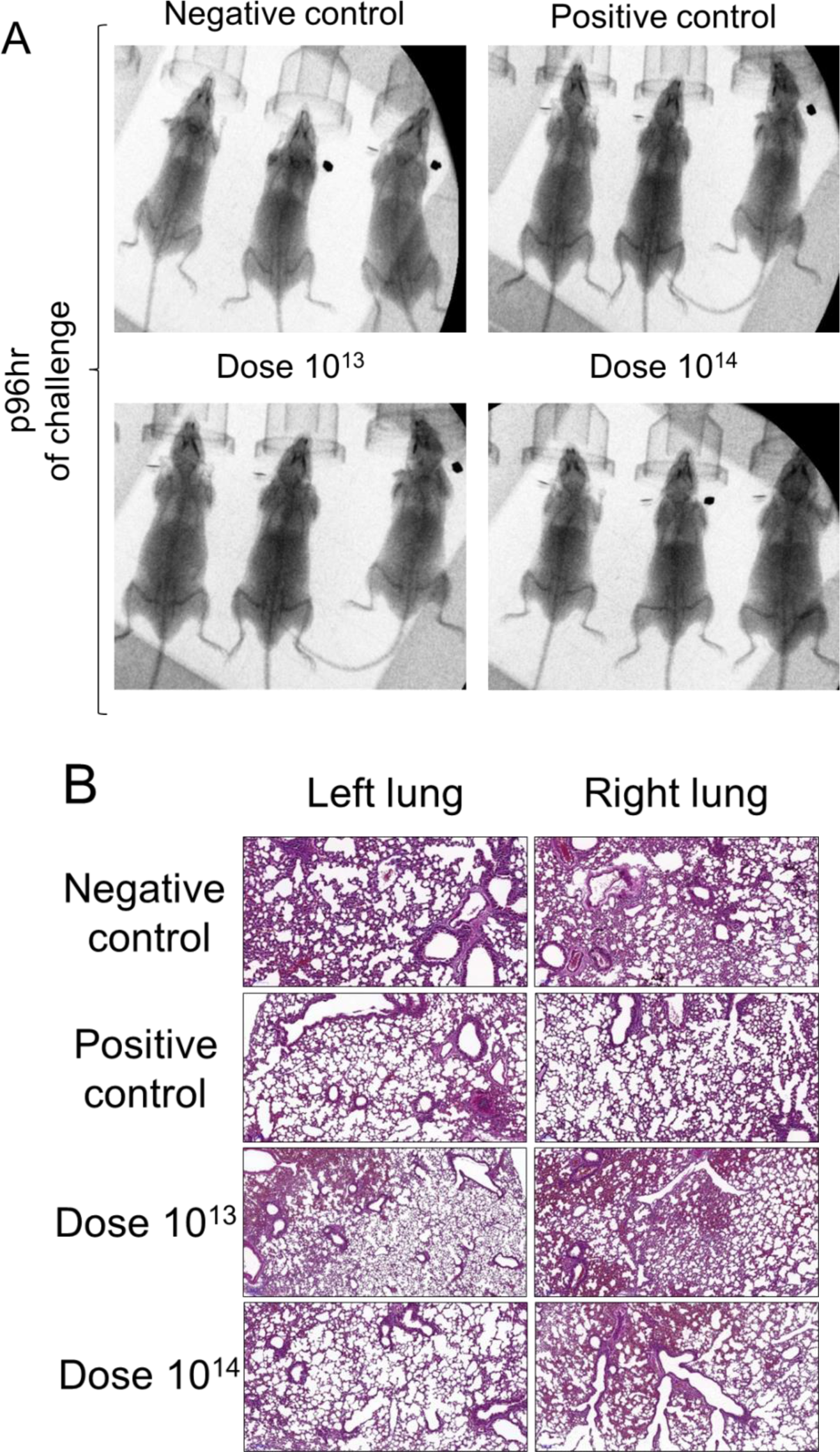
X-ray imaging and histopathology analysis of the Lungs of SARS-CoV-2 infected mice. **A.** X-ray imaging and **B.** Histopathology analysis of the mice groups that were negative control (uninfected and no vaccination), positive control (only infection), dose 10^13^ group, and dose 10^14^ group (vaccinated and infected).

### In vitro efficacy analysis of serum and T cells isolated from mice following challenge test

SARS-CoV-2 specific IgG antibody analysis was performed in 1: 128 and 1: 256 titrations of serum isolated from blood. In SARS-CoV-2 antibody measurements, antibody development was observed in the vaccine groups, including the virus-administered group (**Fig. 5A**). According to the IgG ELISA result, the SARS-CoV-2 IgG antibody increase was significantly detected at 1:256 dilutions in the dose 10^14^ vaccinated group compared to the positive control group (non-vaccinated) (p=0.001) (**Fig. 5A**). The neutralizing antibody study also showed a significant increase in both vaccine groups compared to the positive control at 1: 256, similar to the antibody levels (**Fig. 5B**). However, there was no significant change in gamma interferon responses from mouse spleen T cells without re-stimulation (**Fig. 5C**). Next, we wanted to determine the cytokine secretion profile and T cell frequencies between groups without a re-stimulation. Although it was not statistically significant, TNFα secretion was also seen to increase in the dose 10^14^ group (**Fig. 5D**). The increase of IL-2 in the highest-dose (10^14^) vaccine group indicates that the mice vaccinated after viral challenge show a Th1 type response (**Fig. 5D**). Also, when we compare SARS-CoV-2 infected non-vaccinated positive control with non-vaccinated and un-infected negative control, we determined that IL-10 cytokine, known as cytokine synthesis inhibitory factor, was significantly increased (**Fig. 5D**), suggesting downregulation of the expression of cytokines (Eskdale et al. 1997). On the other hand, we wanted to determine a change in the proportion of spleen T cell subsets upon re-stimulation with SARS-CoV-2 peptides (**Fig. 5E**). Although total CD3+ T and CD4+ T cell populations did not increase in the vaccinated groups regarding control groups, CD25+ CD4+ T cell population was determined to increase in dose groups (**Fig. 5E**). Depending on the viral challenge, frequencies of CD3+ and CD4+ T lymphocytes significantly increased in the positive control group (non-vaccinated viral challenge group), while this increase was not observed in the vaccinated group (**Fig. 5E**). Only an increase in the amount of activated (CD25+) CD4+ T cell was observed in the vaccinated groups (**Fig. 5E**). This data showing that SARS-CoV-2 viral infection was caused to stimulate T cell response along with increase of Th1 inhibitory Tr1 (T cell regulatory)-related IL-10 cytokine secretion and with the absence of Th2-related cytokine response. To sum up, the in vitro efficacy analysis of the challenge test showed that the presence of active T lymphocytes significantly increased in the highest dose (10^14^) vaccine group. The study indicated that viral dissemination was blocked by SARS-CoV-2 specific antibodies and neutralizing antibodies. It was also determined that the ADE effect was not observed, and also confirming that OZG-38.61.3 was non-replicative. As the cellular immune response, CD4+ T cell activation was present, especially at the highest dose, and T cell response was biased to the Th1 response type as desired in the immunization.

**Figure 5:**
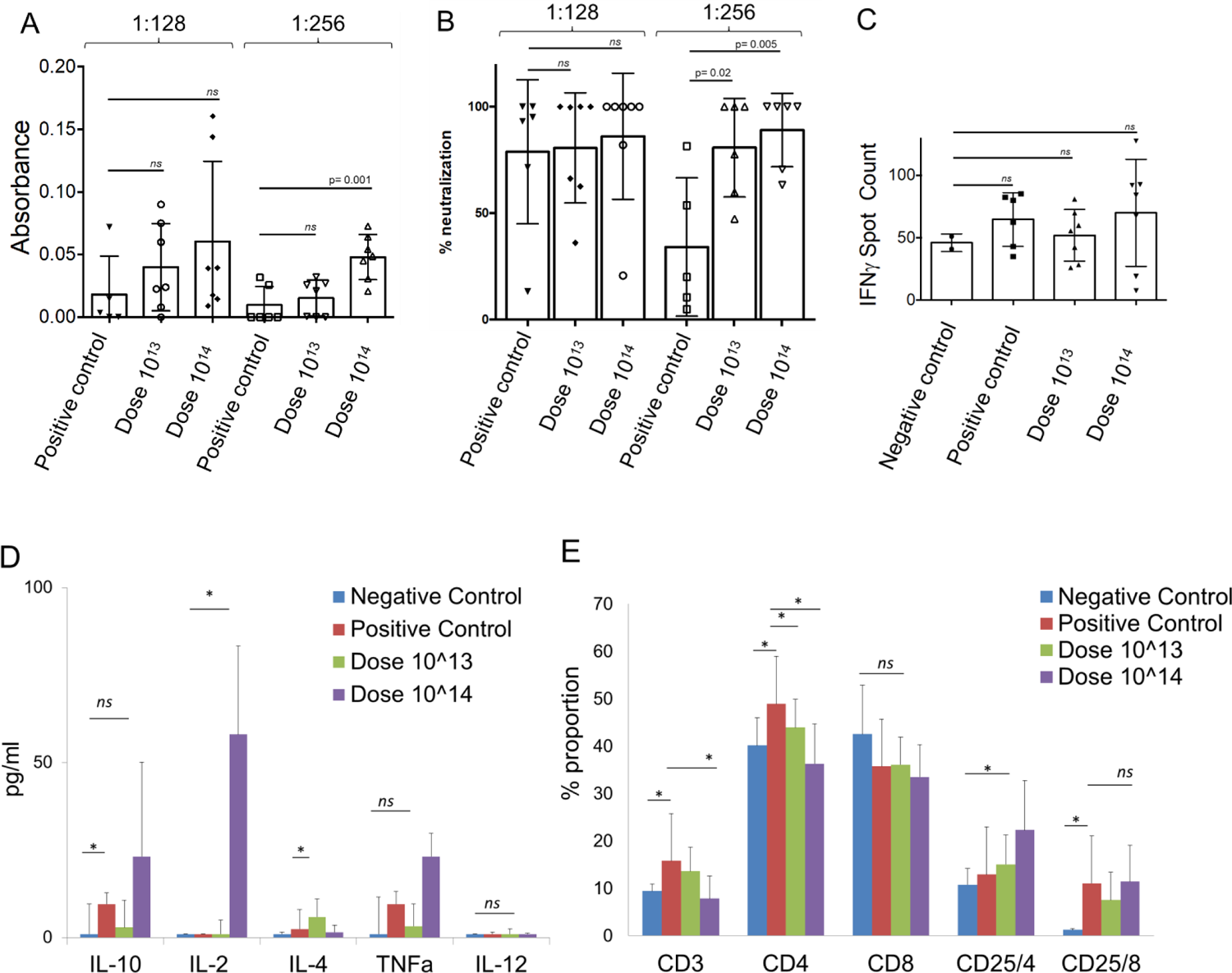
In vitro efficacy analysis of serum and spleen T cell isolated from challenge applied mice. **A.** The bar graph showing presence (absorbance) of SARS-CoV-2 specific IgG in the mice sera diluted with either 1:128 or 1:256 detected using ELISA. **B.** The bar graph showing neutralization frequency of the mice sera diluted to 1:128 or 1:256 that were pre-incubated with a 100x TCID50 dose of infective SARS-CoV-2. The analysis was performed with MTT analysis at 96hr. **C.** The bar graph showing activation frequency (IFNγ positive spot count) of spleen T cells that were incubated with SARS-CoV-2 specific peptides. **D.** The bar graph showing cytokine (IL-10, IL-2, IL-4, TNFa, and IL-12) proportions of spleen T cells that were incubated with SARS-CoV-2 specific peptides. **E.** The bar graph showing a change in the frequency of the T cells after the stimulation. The activation of the T cells was determined with the upregulation of the CD25 surface marker.

## Discussion

The SARS-CoV-2 virus caused one of the severest pandemic around the world. The safe and effective vaccine development for urgent use became more of an issue to end the global COVID-19 pandemic. Several vaccine candidates have recently begun clinical phase studies, and many others are in preclinical development (Gao et al. 2020b; L. Corey et al. 2020). Here, we optimized an inactivated virus vaccine which includes the gamma irradiation process for the inactivation as an alternative to classical chemical inactivation methods so that there is no extra purification required. Previous studies showed that gamma-radiation can induce immunogenicity more effectively rather than conventional inactivation procedures (Seo 2015). Also, we applied the vaccine candidate (OZG-38.61.3) using the intradermal route in mice which decreased the requirement of a higher concentration of inactivated virus for proper immunization unlike most of the classical inactivated vaccine treatments (Lambert and Laurent 2008; Hickling et al. 2011).

Different variations in SARS-CoV-2 strains may occur when producing large quantities (bulk) of the virus in a laboratory (Zhang et al. 2019). For this reason, 50% of the unit volume of virus isolates cultured in multi-layered flasks was frozen in each passage. While preparing the final product (OZG-38.61.3), frozen raw intermediate products were pooled. Thus, pre-pooling genomic characterization of individual variants between passages was made and the final product was created for a more effective and safer vaccine design. At the end of the vaccine production, the final product was found to contain most of the defined mutations in the SARS-CoV-2 strain. In addition, the SARS-CoV-2 virus was passaged 3 times for the isolation from the first donor and 6 times for the final production of OZG-38.61.3. The genome analysis of the OZG-38.61.3 vaccine in this study was found to retain >%99.5 homology with the starting virus stock isolated from the COVID-19 patient. This may enable our inactive virus vaccine to be effective in a large population.

Zeta-sizer along with Nanosight size analysis, proteome, and electron microscopic data showed that the OZG-38.61.3 vaccine preserved its compact structure despite gamma irradiation and lyophilization. However, we also detected aggregate formation, especially in electron microscope images. We added human serum albumin (<0.02%) to the final product to increase the stability, to prevent viral particles from adhering to the injection vial walls, and efficacy of the vaccine candidate (Prymula et al. 2016). Assessment of the residual Vero host cell protein and DNA level in each vaccine dose in this study showed that the protein level was <4ng and DNA was absent in the dose. This showed us that the vaccine production process is efficiently pure from the residual products.

In this study, we generated a prototype gamma-irradiated inactive SARS-CoV-2 vaccine (OZG-38.61.3) and assessed protective efficacy against the intranasal SARS-CoV-2 challenge in transgenic human ACE2 encoding mice. We demonstrate vaccine protection with substantial ∼3 log10 reductions in mean viral loads in dose 10^14^ immunized mice compared with non-vaccinated infected positive control mice. We showed humoral and cellular immune responses against the SARS-CoV-2, including the neutralizing antibodies similar to those shown in Balb/c mice, without substantial toxicity. This study will lead to the initiation of the Phase 1 clinical application of the vaccine for the COVID-19 pandemic.

When we performed the efficacy and safety test of the final product, OZG-38.61.3, vaccine candidate on Balb/ c mice at two different doses (10^13^ and 10^14^), the presence of SARS-CoV-2 specific neutralizing antibodies was significantly detected in the highest-dose vaccination (10^14^). However, at both dose groups, significant IFNγ secretion from the spleen T cells were detected in comparison with the controls, illustrating that cellular immune response developed earlier than the humoral immune response. The fact that the neutralizing test was more accurate than the IgG ELISA analysis may be due to the increased levels of SARS-CoV-2 specific IgA and IgM antibodies (Woo et al. 2004; Demers-Mathieu et al. 2020; Poland, Ovsyannikova, and Kennedy 2020). Moreover, mice vaccinated with both doses showing a significant increase in T cell responses and Th1 dominant cytokine release is an additional evidence of vaccine efficacy. As no significant toxicity was encountered in the histopathological analysis of Balb/c mice vaccinated with both doses, decision was taken to proceed to the challenge test.

A difficulty was faced with the intradermal vaccination of mice in that some of the study mice had skin injury due to the vaccination. This factor may have possibly reduced the efficacy of the intradermal vaccine tests and may be the reason behind the finding of a high standard deviation and inability to see a parallel neutralization capacity in each mouse.

In the Challenge test, we collected oropharyngeal samples to determine the viral copy number following the administration of intranasal infective SARS-CoV-2 virus. In unvaccinated but virus-infected positive control mice viral copy numbers at 48 and 96 hours either did not change or they were increased. However, in the groups of mice vaccinated with both doses, findings showed that copy numbers effectively decreased around 3 log10, and even a few mice had completely lost the viral load. X-rays were performed to search for similar effect in the lung lobes, but the classic COVID-19 infection image was not observed in any of the groups. Also qRT-PCR studies and histopathological lung analyses did not reveal pathological changes in the lung tissues. This may have been either due to the short 96-hour infection period, or low amount of virus (30,000 TCID50) used in the 96-hour challenge test might not have been sufficient to descend into the lungs during this period. There are also previous reports regarding the absence of viral load in the lung tissue being due to the amount of infected dose or the virus could be detected in the specific locations of the lung (Subbarao et al. 2004; Gao et al. 2020b). When neutralizing antibody capacity of mice vaccinated in the Challenge test was analyzed, both doses of vaccination were observed to significantly neutralize the SARS-CoV-2. This is in accordance with the previous finding of reduction in viral copy numbers in immunized mice.

When we looked at the neutralizing antibody capacity of mice vaccinated within the scope of the Challenge test, we observed that both doses of vaccination could significantly neutralize SARS-CoV-2. This shows us that the reduction in viral copy rates is consistent. However, when we looked at the T cell response, we could not see any difference in IFNγ release. Presumably, because groups of mice are infected with the virus, T cells may already be stimulated and this may not make a difference in IFNγ release. A significant decrease in CD3+ and CD4+ T cell ratios and an increase in CD25+ CD4+ T cell ratio show that these cells have already been activated. On the other hand, the fact that the virus was neutralized here prevented the increase in CD3+ T cell proportion, therefore viral challenge resulted in only the increase of active T cell. When we looked at spleen T cells that were not re-stimulated, we detected Th1-type cytokine release, as we expected, especially in the 10^14^ dose vaccine group. On the other hand, the significant increase in the ratios of total CD3+ and CD4+ T cells and the ratios of activated (CD25+) CD8+ T cells and the level of the Th1 cytokine and inhibitor IL-10 between the negative control and positive control mice that had an only viral infection. It shows that in a short time such as 96 hours, it started to generate T cell response more effectively than antibody response. This has shown that T cell response occurs in individuals exposed to the virus without sufficient time for neutralizing antibody formation.

In summary, this study demonstrated that the OZG-38.61.3 vaccine candidates created with gamma-irradiated inactivated SARS-CoV-2 viruses produced neutralizing antibodies, especially effective in the 1014 viral copy formulation, and this was effective in protecting transgenic human ACE2 expressing mice agianst SARS-CoV-2 virus. Vaccine candidates were demonstrated to be safe to the tissues of Balb/ c and transgenic mice. This preclinical study lead to phase 1 vaccine clinical trials of OZG-38.61.3 vaccine for the COVID-19 pandemic.

## Supplementary figures/tables

**Supplementary Figure 1.**
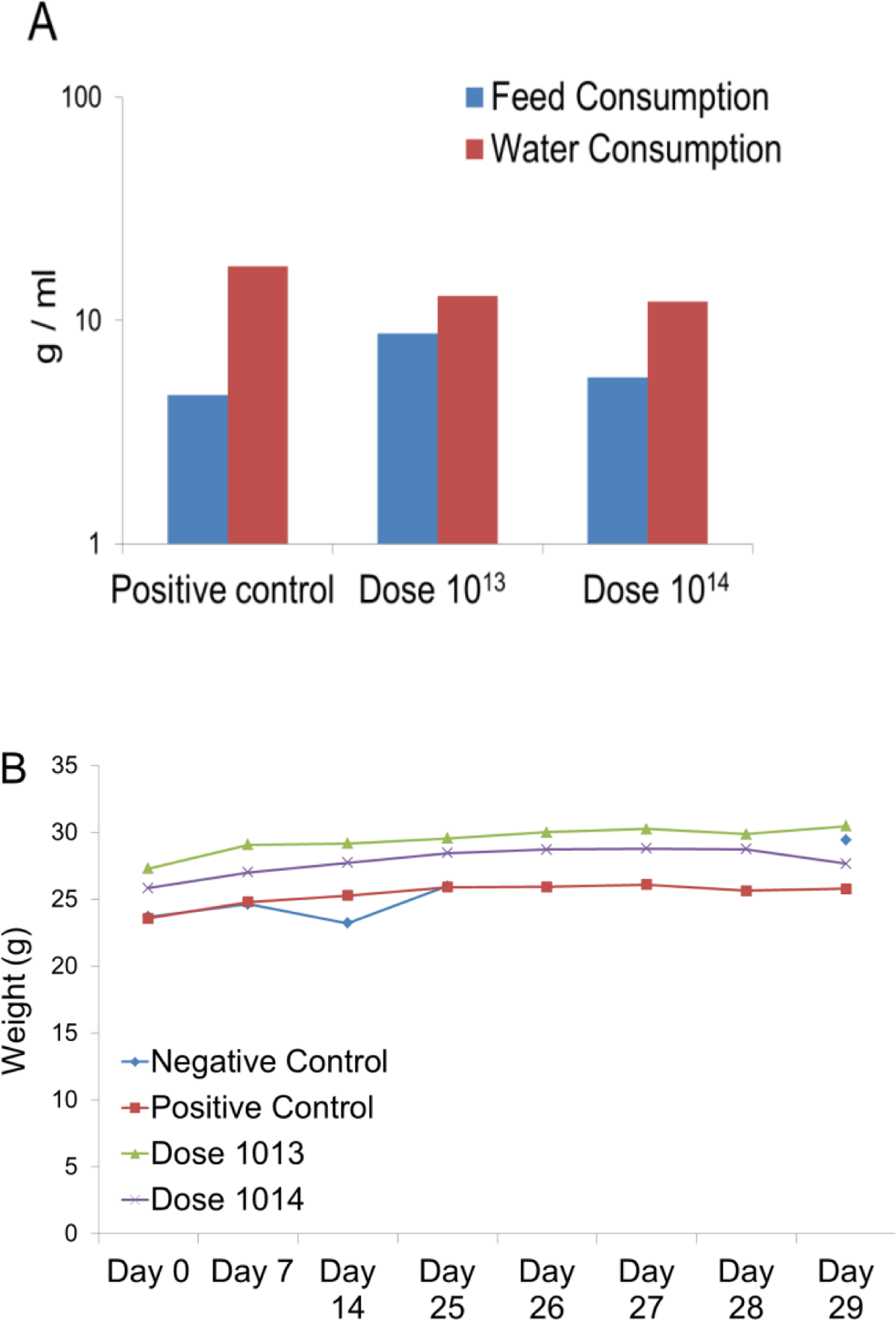
Health state of challenge test mice. **A.** Feed and Water consumption per day post-infection of the positive control group, dose 1×10^13,^ and 1×10^14^ groups. **B.** Monitoring. Body weight changes during challenge experiments.

**Supplementary Table 1:** Final process of lyophilized product of OZG-38.61.3 vaccine candidate. *Infective SARS-CoV-2 viral copy number in a dose.

| Parameter | Method | Limit | Result |
| --- | --- | --- | --- |
| **Appearance** | Visual inspection | Dry Grayish Powder | Passed |
| **Identity** | Single Radial Immunodiffusion (SRID) | Positive | Positive |
|  | TEM | Positive | Positive |
| **Size And Particle Analysis** | Nanosight Zetasizer | Size <200 Nm<br>Particle >1x10<sup>8</sup> MI | Passed |
| **Virus Copy Number/Dose Analysis** | qRT-PCR | *$\geq 1 \times 10^{13}$<br>*$\geq 1 \times 10^{14}$ | Passed |
| **Biochemical Analysis** | Ph, Na, Cl, K, Ca, Glucose, Total Protein, Albumin | Reference Interval<br>$\geq 70 \mu\text{g}$ total protein/ dose | Passed |
| **Relative Humidity** | Moisture Analyzer | <%3 | Passed |
| **Purity** | Vero Protein ELISA | <4 ng | Passed |
|  | Vero Dna Nanodrop | <10 ng | Passed |
| **Replicative Virus Test** | Real-Time Cell Analysis (RTCA) or MTT | - | Passed |
| **Proteome** | Mass Spectrometer | - | Passed |
| **Microbiological Quality Control** | Culture | Negative | Negative |
|  | Gram Stain | Negative | Negative |
| **Fungi** | Culture | Negative | Negative |
| **Mycoplasma** | PCR | Negative | Negative |
| **Endotoxin** | Lal Test | <0.5 IU/ml | Passed |
| **Stability/ Efficiency** | Single Radial Immunodiffusion (SRID) | Positive | Efficiency analysis of 0,1,6,12,24 and 26 months in 4-24°C storage conditions. |

**Supplementary Table 2:**
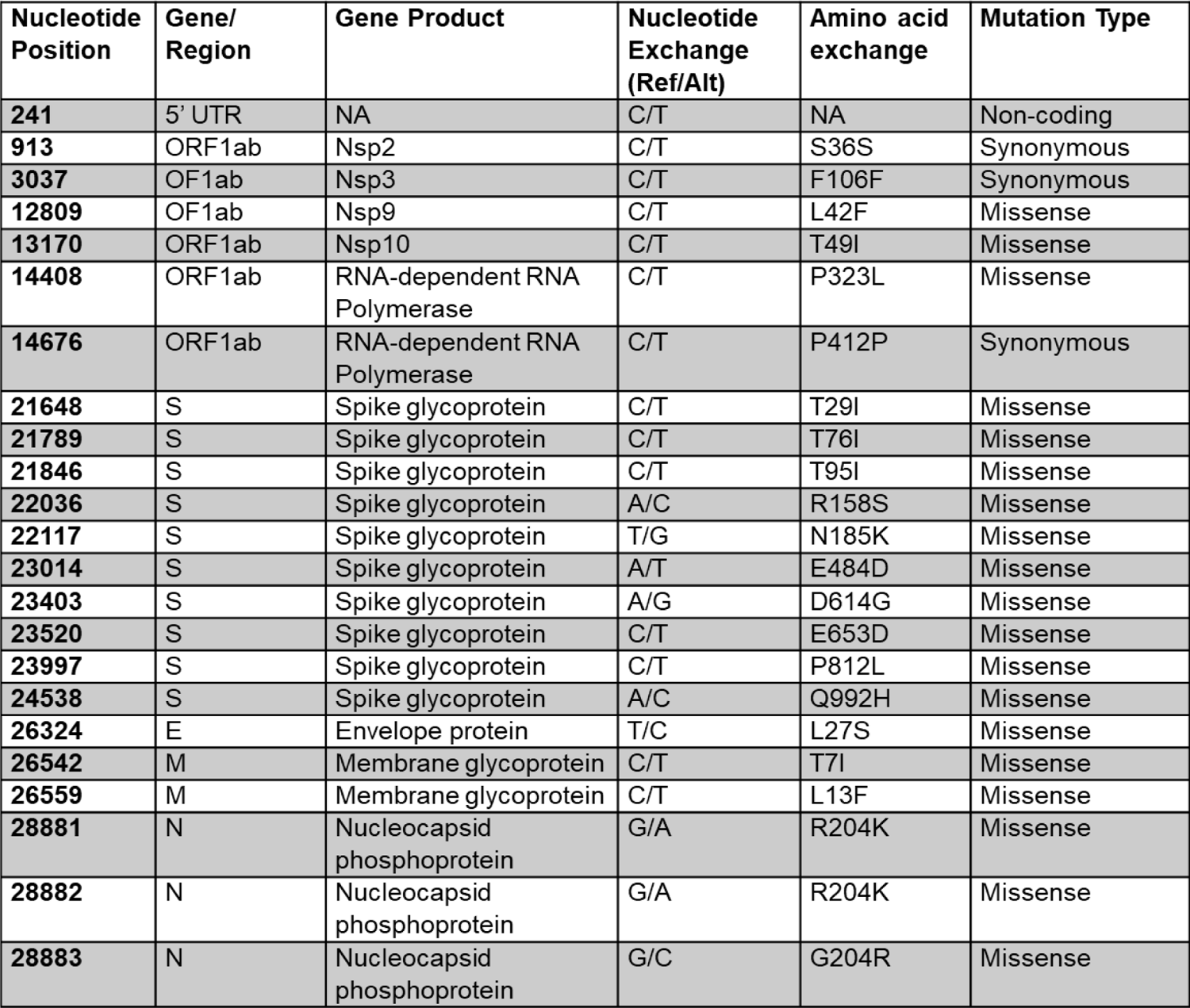
Viral Genome Sequencing analysis of fractionated SARS-CoV-2 strains forming OZG-38.61.3 vaccine candidate.

| Nucleotide Position | Gene/ Region | Gene Product | Nucleotide Exchange (Ref/Alt) | Amino acid exchange | Mutation Type |
| --- | --- | --- | --- | --- | --- |
| 241 | 5' UTR | NA | C/T | NA | Non-coding |
| 913 | ORF1ab | Nsp2 | C/T | S36S | Synonymous |
| 3037 | OF1ab | Nsp3 | C/T | F106F | Synonymous |
| 12809 | OF1ab | Nsp9 | C/T | L42F | Missense |
| 13170 | ORF1ab | Nsp10 | C/T | T49I | Missense |
| 14408 | ORF1ab | RNA-dependent RNA Polymerase | C/T | P323L | Missense |
| 14676 | ORF1ab | RNA-dependent RNA Polymerase | C/T | P412P | Synonymous |
| 21648 | S | Spike glycoprotein | C/T | T29I | Missense |
| 21789 | S | Spike glycoprotein | C/T | T76I | Missense |
| 21846 | S | Spike glycoprotein | C/T | T95I | Missense |
| 22036 | S | Spike glycoprotein | A/C | R158S | Missense |
| 22117 | S | Spike glycoprotein | T/G | N185K | Missense |
| 23014 | S | Spike glycoprotein | A/T | E484D | Missense |
| 23403 | S | Spike glycoprotein | A/G | D614G | Missense |
| 23520 | S | Spike glycoprotein | C/T | E653D | Missense |
| 23997 | S | Spike glycoprotein | C/T | P812L | Missense |
| 24538 | S | Spike glycoprotein | A/C | Q992H | Missense |
| 26324 | E | Envelope protein | T/C | L27S | Missense |
| 26542 | M | Membrane glycoprotein | C/T | T7I | Missense |
| 26559 | M | Membrane glycoprotein | C/T | L13F | Missense |
| 28881 | N | Nucleocapsid phosphoprotein | G/A | R204K | Missense |
| 28882 | N | Nucleocapsid phosphoprotein | G/A | R204K | Missense |
| 28883 | N | Nucleocapsid phosphoprotein | G/C | G204R | Missense |

**Supplementary Table 3:** LC-MS/MS results of candidate vaccine sample developed against the SARS-CoV-2 virus. OX=2697049.

| Accession | Description | Coverage [%] | # PS M | # Peptide | # AAs | MW [kDa] | calc. pI | Score Sequences HT |
| --- | --- | --- | --- | --- | --- | --- | --- | --- |
| A0A6G7SLW8 | Orf1ab polyprotein OS=Severe acute respiratory syndrome coronavirus 2 | 1,02860 | 19 | 4 | 7097 | 793,7 | 6,76 | 0 |
| A0A6C0T6Z7 | Nucleoprotein OS=Severe acute respiratory syndrome coronavirus 2 O | 42,0047 | 30 | 12 | 419 | 45,6 | 10,07 | 56,69 |
| A0A679G9E9 | S protein OS=Severe acute respiratory syndrome coronavirus 2 | 0,54988 | 1 | 1 | 1273 | 141,1 | 6,65 | 0 |
| A0A6C0N6C6 | Nonstructural protein NS8 OS=Severe acute respiratory syndrome coronavirus2 | 23,9669 | 2 | 2 | 121 | 13,8 | 5,73 | 0 |
| A0A6H2LA46 | ORF1ab polyprotein OS=Severe acute respiratory syndrome coronavirus 2 | 1,02874 | 19 | 4 | 7096 | 792,5 | 6,7 | 0 |
| A0A6H2LAX8 | ORF1ab polyprotein OS=Severe acute respiratory syndrome coronavirus 2 | 1,02874 | 19 | 4 | 7096 | 793,5 | 6,76 | 0 |
| A0A6H1PMM5 | Surface glycoprotein OS=Severe acute respiratory syndrome coronavirus 2 | 0,54988 | 1 | 1 | 1273 | 141,1 | 6,61 | 0 |
| A0A6H2L905 | ORF1ab polyprotein OS=Severe acute respiratory syndrome coronavirus 2 | 1,02874 | 19 | 4 | 7096 | 793,5 | 6,76 | 0 |
| A0A6H2LC46 | Surface glycoprotein OS=Severe acute respiratory syndrome coronavirus 2 | 0,54988 | 1 | 1 | 1273 | 140,3 | 6,76 | 0 |
| A0A6H2LCV0 | ORF1ab polyprotein OS=Severe acute respiratory syndrome coronavirus 2 | 1,02874 | 19 | 4 | 7096 | 793,5 | 6,76 | 0 |
| A0A6H2L673 | Surface glycoprotein OS=Severe acute respiratory syndrome coronavirus 2 | 0,54988 | 1 | 1 | 1273 | 141,1 | 6,73 | 0 |
| A0A6H2L7S6 | ORF1ab polyprotein OS=Severe acute respiratory syndrome coronavirus 2 | 1,02874 | 19 | 4 | 7096 | 793,7 | 6,76 | 0 |
| A0A6H2L8K8 | ORF1ab polyprotein OS=Severe acute respiratory syndrome coronavirus 2 | 1,02874 | 19 | 4 | 7096 | 793,5 | 6,62 | 0 |
| A0A6H2L9B8 | ORF1ab polyprotein OS=Severe acute respiratory syndrome coronavirus 2 | 1,02874 | 19 | 4 | 7096 | 793,6 | 6,76 | 0 |
| A0A6H2LBL6 | ORF1ab polyprotein OS=Severe acute respiratory syndrome coronavirus 2 | 1,02874 | 19 | 4 | 7096 | 793,6 | 6,76 | 0 |
| A0A6H2LDA7 | ORF1ab polyprotein OS=Severe acute respiratory syndrome coronavirus 2 | 1,02874 | 19 | 4 | 7096 | 792,9 | 6,64 | 0 |
| A0A6H2LBC7 | ORF1ab polyprotein OS=Severe acute respiratory syndrome coronavirus 2 | 1,02874 | 19 | 4 | 7096 | 793,2 | 6,68 | 0 |
| A0A6H2LA93 | Nucleocapsid phosphoprotein OS=Severe acute respiratory syndrome coronavirus 2 | 42,0047 | 30 | 12 | 419 | 45,6 | 10,07 | 56,69 |
| A0A6H2L973 | ORF1ab polyprotein OS=Severe acute respiratory syndrome coronavirus 2 | 1,02874 | 19 | 4 | 7096 | 793 | 6,81 | 0 |
| A0A6H2L8H9 | ORF1ab polyprotein OS=Severe acute respiratory syndrome coronavirus 2 | 1,02874 | 19 | 4 | 7096 | 793,6 | 6,76 | 0 |
| A0A6H2LAC5 | ORF1ab polyprotein OS=Severe acute respiratory syndrome coronavirus 2 | 1,02874 | 19 | 4 | 7096 | 793,3 | 6,71 | 0 |
| A0A6H2LBD2 | Surface glycoprotein OS=Severe acute respiratory syndrome coronavirus 2 | 0,54988 | 1 | 1 | 1273 | 141 | 6,73 | 0 |
| A0A6H2L6R1 | ORF1ab polyprotein (Fragment) OS=Severe acute respiratory syndrome coronavirus 2 | 1,04539 | 19 | 4 | 6983 | 780,6 | 6,77 | 0 |
| A0A6H2L7I9 | ORF1ab polyprotein OS=Severe acute respiratory syndrome coronavirus 2 | 1,02874 | 19 | 4 | 7096 | 793,2 | 6,7 | 0 |
| A0A6H2LC40 | Surface glycoprotein OS=Severe acute respiratory syndrome coronavirus 2 | 0,54988 | 1 | 1 | 1273 | 140,9 | 6,73 | 0 |
| A0A6H2LBU1 | Surface glycoprotein (Fragment) OS=Severe acute respiratory syndrome coronavirus 2 | 0,61946 | 1 | 1 | 1130 | 124,9 | 6,44 | 0 |
| A0A6H2LC49 | ORF1ab polyprotein OS=Severe acute respiratory syndrome coronavirus 2 | 1,02874 | 19 | 4 | 7096 | 793,6 | 6,76 | 0 |
| A0A6H2L683 | Surface glycoprotein OS=Severe acute respiratory syndrome coronavirus 2 | 0,54988 | 1 | 1 | 1273 | 141,1 | 6,81 | 0 |
| A0A6H2LDI3 | ORF1ab polyprotein OS=Severe acute respiratory syndrome coronavirus 2 | 1,02874 | 19 | 4 | 7096 | 793,5 | 6,73 | 0 |
| A0A6H2L877 | ORF1ab polyprotein OS=Severe acute respiratory syndrome coronavirus 2 | 1,02874 | 19 | 4 | 7096 | 793,6 | 6,76 | 0 |
| **A0A6H2L915** | Nucleocapsid phosphoprotein OS=Severe acute respiratory syndrome coronavirus 2 | 42,0047 | 30 | 12 | 419 | 45,7 | 10,05 | 56,69 |
| **A0A6H2LBN3** | ORF8 protein OS=Severe acute respiratory syndrome coronavirus 2 | 23,9669 | 2 | 2 | 121 | 13,8 | 5,73 | 0 |
| **A0A6H2L880** | ORF1ab polyprotein OS=Severe acute respiratory syndrome coronavirus 2 | 1,02874 | 19 | 4 | 7096 | 793,1 | 6,62 | 0 |
| **A0A6H2LAL1** | ORF1ab polyprotein OS=Severe acute respiratory syndrome coronavirus 2 | 1,02874 | 19 | 4 | 7096 | 793,6 | 6,76 | 0 |
| **A0A6H2L964** | Surface glycoprotein OS=Severe acute respiratory syndrome coronavirus 2 | 0,54988 | 1 | 1 | 1273 | 141,1 | 6,73 | 0 |
| **A0A6H2L9W1** | ORF1ab polyprotein OS=Severe acute respiratory syndrome coronavirus 2 | 1,02874 | 19 | 4 | 7096 | 793,6 | 6,76 | 0 |
| **A0A6H2LCH0** | ORF1ab polyprotein OS=Severe acute respiratory syndrome coronavirus 2 | 1,02874 | 19 | 4 | 7096 | 793,6 | 6,76 | 0 |
| **A0A6H2L702** | Surface glycoprotein OS=Severe acute respiratory syndrome coronavirus 2 | 0,54988 | 1 | 1 | 1273 | 141,1 | 6,65 | 0 |
| **A0A6H1PRW5** | ORF1ab polyprotein OS=Severe acute respiratory syndrome coronavirus 2 | 1,02874 | 19 | 4 | 7096 | 793,6 | 6,76 | 0 |
| **A0A6H2L7Z5** | Nucleocapsid phosphoprotein OS=Severe acute respiratory syndrome coronavirus 2 | 42,0047 | 30 | 12 | 419 | 45,6 | 10,07 | 56,69 |
| **A0A6H2L6T0** | Nucleocapsid phosphoprotein OS=Severe acute respiratory syndrome coronavirus 2 | 42,0047 | 30 | 12 | 419 | 45,6 | 10,07 | 56,69 |
| **A0A6H2LA09** | ORF1ab polyprotein OS=Severe acute respiratory syndrome coronavirus 2 | 1,02874 | 19 | 4 | 7096 | 793 | 6,71 | 0 |
| **A0A6H2LCI8** | ORF1ab polyprotein OS=Severe acute respiratory syndrome coronavirus 2 | 1,02874 | 19 | 4 | 7096 | 793,6 | 6,76 | 0 |
| **A0A6H2L7Z4** | ORF1ab polyprotein OS=Severe acute respiratory syndrome coronavirus 2 | 1,02874 | 19 | 4 | 7096 | 793,7 | 6,76 | 0 |
| **A0A6H2LA55** | ORF1ab polyprotein OS=Severe acute respiratory syndrome coronavirus 2 | 1,02874 | 19 | 4 | 7096 | 793,6 | 6,8 | 0 |
| **A0A6H2L7V1** | Nucleocapsid phosphoprotein OS=Severe acute respiratory syndrome coronavirus 2 | 42,0047 | 30 | 12 | 419 | 45,6 | 10,07 | 56,69 |
| **A0A6H2LD79** | ORF1ab polyprotein OS=Severe acute respiratory syndrome coronavirus 2 | 1,02874 | 19 | 4 | 7096 | 793,6 | 6,76 | 0 |
| **A0A6H2L9L2** | ORF1ab polyprotein OS=Severe acute respiratory syndrome coronavirus 2 | 1,02874 | 19 | 4 | 7096 | 793,2 | 6,71 | 0 |
| **A0A6H2L6I9** | ORF1ab polyprotein OS=Severe acute respiratory syndrome coronavirus 2 | 1,02874 | 19 | 4 | 7096 | 792,9 | 6,7 | 0 |
| **A0A6H2LAG5** | Surface glycoprotein OS=Severe acute respiratory syndrome coronavirus 2 | 0,54988 | 1 | 1 | 1273 | 140,6 | 6,04 | 0 |
| **A0A6H2L8N1** | ORF1ab polyprotein OS=Severe acute respiratory syndrome coronavirus 2 | 1,02874 | 19 | 4 | 7096 | 793,6 | 6,76 | 0 |
| **A0A6G7SLV1** | Orf1ab polyprotein OS=Severe acute respiratory syndrome coronavirus 2 | 1,02874 | 19 | 4 | 7096 | 793,6 | 6,76 | 0 |
| **A0A6H2EH11** | ORF1ab polyprotein OS=Severe acute respiratory syndrome coronavirus 2 | 1,02874 | 19 | 4 | 7096 | 793,5 | 6,76 | 0 |
| **A0A6H0QT63** | ORF1ab polyprotein OS=Severe acute respiratory syndrome coronavirus 2 | 1,02874 | 19 | 4 | 7096 | 793,5 | 6,76 | 0 |
| **A0A6H2L7L3** | Surface glycoprotein OS=Severe acute respiratory syndrome coronavirus 2 | 0,54988 | 1 | 1 | 1273 | 141 | 6,81 | 0 |
| **A0A6H2L5M8** | Nucleocapsid phosphoprotein OS=Severe acute respiratory syndrome coronavirus 2 | 42,0047 | 30 | 12 | 419 | 45,7 | 10,08 | 56,69 |
| **A0A6H2LAI7** | ORF1ab polyprotein OS=Severe acute respiratory syndrome coronavirus 2 | 1,02874 | 19 | 4 | 7096 | 793,6 | 6,76 | 0 |
| **A0A6H2LEE1** | ORF1ab polyprotein OS=Severe acute respiratory syndrome coronavirus 2 | 1,02874 | 19 | 4 | 7096 | 793,6 | 6,76 | 0 |
| **A0A6H2LBP4** | ORF1ab polyprotein OS=Severe acute respiratory syndrome coronavirus 2 | 1,02874 | 19 | 4 | 7096 | 793,6 | 6,74 | 0 |
| **A0A6H2L794** | ORF1ab polyprotein OS=Severe acute respiratory syndrome coronavirus 2 | 1,02874 | 19 | 4 | 7096 | 793,6 | 6,76 | 0 |
| **A0A6H2LCV8** | ORF1ab polyprotein OS=Severe acute respiratory syndrome coronavirus 2 | 1,02874 | 19 | 4 | 7096 | 793,6 | 6,76 | 0 |
| **A0A6C0RS15** | 2'-O-methyltransferase OS=Severe acute respiratory syndrome coronavirus2 | 1,028749 | 19 | 4 | 7096 | 793,6 | 6,76 | 0 |
| **A0A6G7SLW0** | Orf1ab polyprotein OS=Severe acute respiratory syndrome coronavirus 2 | 1,02874 | 19 | 4 | 7096 | 793,5 | 6,74 | 0 |
| **A0A6H2LB79** | ORF1ab polyprotein OS=Severe acute respiratory syndrome coronavirus 2 | 1,028749 | 19 | 4 | 7096 | 793,6 | 6,76 | 0 |
| **A0A6H2LB01** | ORF1ab polyprotein OS=Severe acute respiratory syndrome coronavirus 2 | 1,02874 | 19 | 4 | 7096 | 792,8 | 6,71 | 0 |

**Supplementary Table 4:** **Organ weights and histopathological analysis vaccinated Balb/c mice. A.** Organ Weights Chart. Histological Analysis of **B.** the lung. **C.** the kidney **D.** the liver tissues. Control mice, n=5, Dose 10^13^ vaccinated mice, n=5, and Dose 10^14^ vaccinated mice, n=5. **\***mild inflammation

|  | **Control** | **1x10<sup>13</sup>** | **1x10<sup>14</sup>** |
| --- | --- | --- | --- |
| **Total Body Weight** | 35,16±0,9 | 35,16±0,8 | 35,16±0,7 |
| **Lung** | 1,34±0,01 | 1,34±0,03 | 1,343±0,02 |
| **Liver** | 2,59±0,14 | 2,59±0,19 | 2,59±0,18 |
| **Spleen** | 1,22±0,01 | 1,22±0,01 | 1,22±0,01 |
| **Kidney** | 2,64±0,03 | 2,64±0,04 | 2,64±0,05 |

|  | **Control** | **1x10<sup>13</sup>** | **1x10<sup>14</sup>** |
| --- | --- | --- | --- |
| **Inflammation** | 2/5* | 1/5* | 1/5* |
| **Hemorrhage** | 0/5 | 0/5 | 0/5 |
| **Eosinophil** | 0/5 | 1/5 | 0/5 |

|  | **Control** | **1x10<sup>13</sup>** | **1x10<sup>14</sup>** |
| --- | --- | --- | --- |
| **Inflammation** | 0/5 | 2/5* | 0/5 |
| **Hemorrhage** | 0/5 | 0/5 | 0/5 |
| **Eosinophil** | 0/5 | 0/5 | 0/5 |

|  | **Control** | **1x10<sup>13</sup>** | **1x10<sup>14</sup>** |
| --- | --- | --- | --- |
| **Inflammation** | 1/5* | 1/5* | 0/5 |
| **Hemorrhage** | 0/5 | 0/5 | 0/5 |
| **Eosinophil** | 0/5 | 0/5 | 0/5 |

